# Development of a yeast-based vaccine for *Theileria parva* infection in cattle

**DOI:** 10.1101/2021.01.30.428938

**Authors:** Shan Goh, Jeannine Kolakowski, Angela Holder, Mark Pfuhl, Daniel Ngugi, Keith Ballingall, Kata Tombacz, Dirk Werling

## Abstract

East Coast Fever (ECF), caused by the tick-borne apicomplexan parasite *Theileria parva,* remains one of the most important livestock diseases in sub-Saharan Africa with more than 1 million cattle dying from infection every year. Disease prevention relies on the so-called “Infection and Treatment Method” (ITM), which is costly, complex, laborious, difficult to standardise on a commercial scale and results in a parasite strain specific, MHC class I restricted cytotoxic T cell response. We therefore attempted to develop a safe, affordable, stable, orally applicable and potent subunit vaccine for ECF using five different *T. parva* schizont antigens (Tp1, Tp2, Tp9, Tp10 and N36) and *Saccharomyces cerevisiae* as an expression platform. Full-length native Tp2 and Tp9 as well as fragments of native Tp1 were successfully expressed on the surface of *S. cerevisiae*. *In vitro* analyses highlighted that recombinant yeast expressing Tp2 can elicit IFNy responses using PBMCs from ITM-animals, while Tp2 and Tp9 induced IFNy responses from enriched bovine CD8^+^ T cells. A subsequent *in vivo* study showed that oral administration of heat-inactivated, freeze-dried yeast stably expressing Tp2 increased total murine serum IgG over time, but more importantly, induce Tp2 specific serum IgG antibodies in individual mice compared to the control group. While these results will require subsequent experiments to verify induction of protection in neonatal calves, our data indicates that oral application of yeast expressing Theileria antigens could provide an affordable and easy vaccination platform for sub-Saharan Africa. Evaluation of antigen specific cellular immune responses, especially cytotoxic CD8^+^ T cell immunity in cows will further contribute to the development of a yeast-based vaccine for ECF.

## Introduction

East Coast Fever (ECF), caused by the tick-borne apicomplexan parasite *Theileria parva,* remains one of the most important livestock diseases in sub-Saharan Africa. More than one million cattle succumb to ECF every year resulting in economic losses for mostly smallholder farmers of approximately 300 million US dollars^1,2^.

Treatment of ECF with the antiprotozoal drug buparvaquone is expensive and often ineffective in altering disease progression^1,2^. Disease prevention relies on the “Infection and Treatment Method” (ITM). This involves inoculation of three different strains of live *T. parva* sporozoites (Muguga cocktail) and simultaneous application of long-lasting oxytetracycline. ITM production is costly, complex, laborious^3,4^ and difficult to standardise on a commercial scale as each batch varies in parasite number and antigenic diversity^5–7^. Consequently, antigen characterisation of each batch by comparison of genetic markers is necessary for validation of the vaccine^8,9^. Although ITM can induce long-lasting protection, the resulting immunity is strain specific. African buffalo-derived strains pose a high risk of reinfection due to extensive antigenic variability in field strains^10^. Cattle immunised by the ITM can remain carriers of *T. parva* for many years^11,12^ and a source of infectious ticks, raising concerns regarding the introduction of new parasite strains into areas previously free of them^11,13,14^.

For these reasons, the development of subunit vaccines for ECF is an area of intense research. However, a major hurdle in the development of subunit vaccines is the identification of antigens that provide broad protection. In the case of ECF, such immune protection is provided by induction of antigen specific, MHC class I restricted cytotoxic CD8^+^ T cells^15^ and supported by cytokines from and close contact to antigen specific CD4^+^ T helper cells^16^. Several studies report variants in known *T. parva* antigens that are circulating in cattle or buffalo in different regions^9,17–19^. A comprehensive analysis of antigenic variability in cattle-derived *T. parva* was recently reported and the *T. parva* antigens Tp1, Tp2, Tp9, which are potential vaccine candidates, were found to have extensive sequence diversity compared to Tp3, Tp4, Tp5, Tp6, Tp7, Tp8 and Tp10^19^. Buffalo-derived *T. parva* similarly showed extensive antigenic variability in Tp1 and Tp2^9^. In contrast, the widely used Muguga ITM cocktail comprises three strains that are quite similar antigenically^20^. Subunit vaccines offer the opportunity to include a greater repertoire of antigens, particularly with the advances made in the identification of antigenic variants. Factors contributing to antigenic variability are sexual recombination after cross-infection of a single host with multiple *T. parva* strains and, to a lesser extent, genetic drift within defined geographical locations^19,21^.

*Saccharomyces cerevisiae* is an ideal platform for expression of heterologous eukaryote proteins. It is safe to consume and survives in the gastro-intestinal environment^22,23^. Easy genetic manipulation and large-scale production, strong adjuvant properties^24,25^, and long-term antigen stability at room temperature^26^ are other reasons for its suitability as a vaccine platform. Yeast-based vaccines have already been shown to stimulate protective immune responses towards a broad range of bacteria, viruses and parasites *in vivo*^27–30^. Yeast cells are avidly internalised by dendritic cells (DCs) which subsequently mature into potent antigen presenting cells (APCs)^24,31^. Recombinant antigens expressed by such yeasts are delivered into both MHC class I and II pathways and efficiently presented to MHC class II restricted CD4^+^ T helper and MHC class I restricted, CD8^+^ cytotoxic T cells^24,32^. DC production of proinflammatory cytokines such as IL-12 leads to the induction of CTL mediated immunity which makes yeast particularly attractive in the control of East Coast Fever in cattle^15^. This study was designed to develop a novel yeast-based vaccine for protection against *T. parva* infection in cattle using five of the above-mentioned vaccine antigens, including Tp1, Tp2, Tp9, Tp10 and N36. These were chosen based on the number of CTL epitopes, sequence diversity and presentation by MHC class I molecules.

## Materials and Methods

### *E. coli* strains, yeast strains and media

*Escherichia coli* DH5α (Invitrogen), DH10β (New England Biolabs) and TOP10 (Invitrogen) were used for cloning of recombinant plasmids. Positive transformants were selected on LB medium (1% tryptone, 0.5% yeast extract, 1% NaCl; Oxoid) supplemented with ampicillin (100 µg mL^−1^, Sigma Aldrich) for 24h at 37°C. *Saccharomyces cerevisiae* strains EBY100 (MATα AGA1::GAL1-AGA1::URA3 ura3-52 trp1 leu2Δ200 his3Δ200 pep4::HIS3 prb11.6R can1 GAL) (ATCC) and INVSc1 (MATα his3Δ1 leu2 trp1-289 ura3-52 MAT his3Δ1 leu2 trp1-289 ura3-52) (Invitrogen) were routinely grown in YPD medium (1% yeast extract, 2% peptone, 2% dextrose) and used to express recombinant proteins. EBY100 transformants were grown in minimum dextrose agar (0.67% Yeast Nitrogen Base (YNB), 2% glucose, 0.01% leucine) lacking tryptophan, while INVSc1 transformants were grown in SC minimal medium (0.67% YNB, 2% raffinose, 0.01% each of adenine, arginine, cysteine, leucine, lysine, threonine, tryptophan), 0.005% each of aspartic acid, histidine, isoleucine, methionine, phenylalanine, proline, serine, tyrosine, valine) lacking uracil (SC-U) for three days at 30°C. To induce expression of recombinant protein, EBY100 clones were first grown in YNB-CAA medium (0.67% YNB, 0.5% casamino acids) with 2% glucose overnight at 30°C to OD600 of 2-5, then cells were resuspended in YNB-CAA medium with 2% galactose to OD600 of 0.5-1 and grown at 22°C for up to 48h. To induce expression of recombinant protein in INVSc1, cultures were first grown in SC-U overnight at 30°C to OD600 of 5-7, then resuspended in SC-U medium with 2% galactose to OD600 of 0.4 and grown at 30°C for up to 24h. Liquid cultures were agitated at 250 rpm.

### Construction of recombinant plasmids

*T. parva* genes encoding the predicted antigens Tp1, Tp2 and Tp9 were expressed for either surface display in the pYD1 vector (Yeast Display Vector kit, Invitrogen) or for intracellular expression using the pYES2/NTC vector (N-terminal Xpress and C-terminal V5 epitope, Invitrogen). The pYD1 system used the yeast internal Apa1/Ap2 system, which links the Apa2 protein via two disulphide bonds to the Apa1 proteins expressed on the yeast surface. For each gene, the wild-type (WT) form or a codon-optimized version for subsequent expression in the yeast system or in eukaryotic cells were amplified using specified primers (Table 1).

**Table 1.**
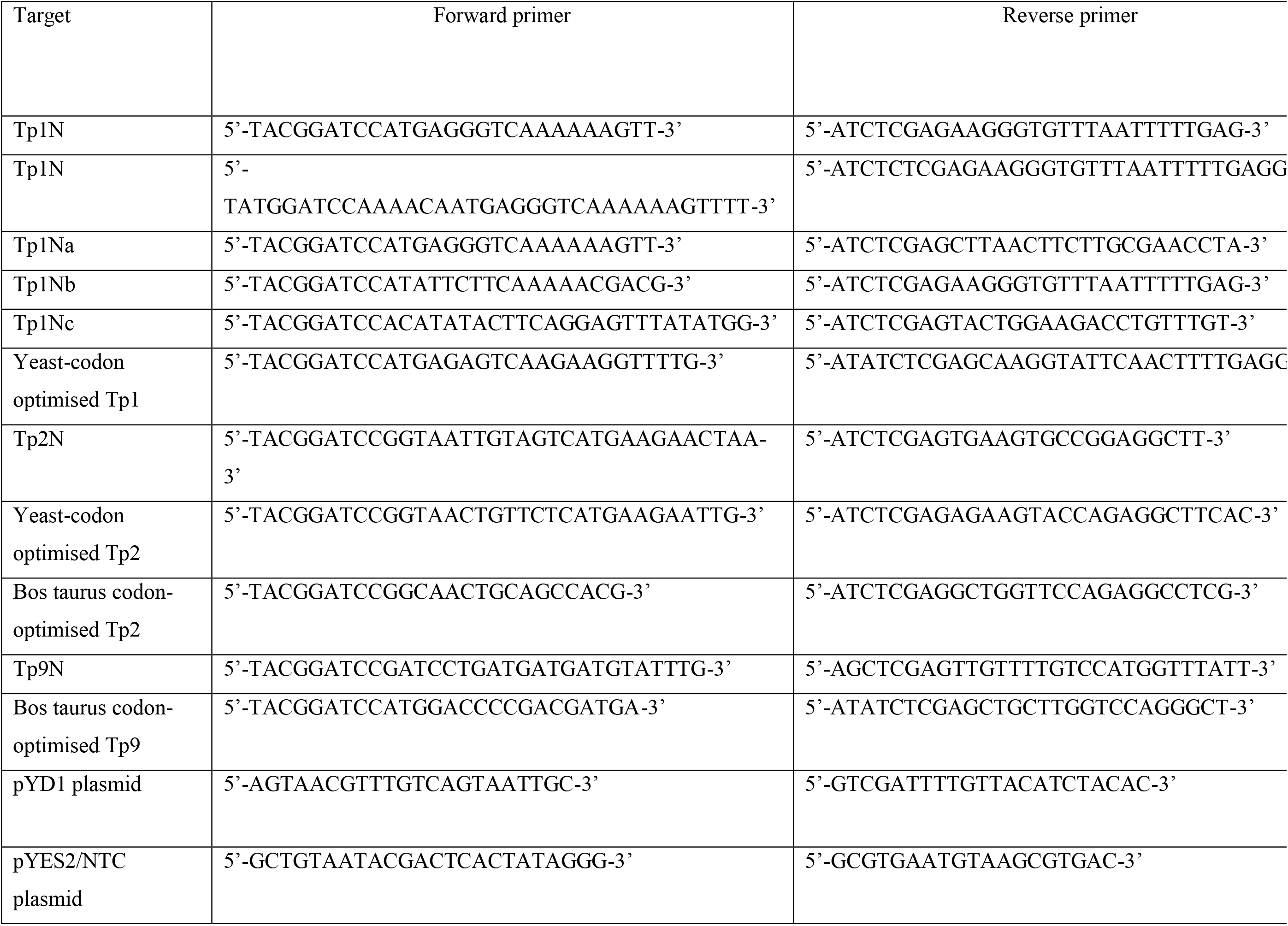
Summary of primers used for cloning of *T. parva* antigen sequences into the pYD1 and pYES2/NTC expression vector

WT gene sequences of Tp1 and Tp2 were amplified from cDNA prepared from total RNA isolated from *T. parva* Muguga infected bovine peripheral blood mononuclear cells (PBMCs) using the Qiagen RNeasy Kit. Contaminating DNA was removed with DNase I (DNA-free removal kit, Invitrogen) and cDNA was prepared with AMV First Strand cDNA kit (New England Biolabs) using oligo-dT to prime the reaction. PCR was carried out with gene-specific primers (Table 1) using Phusion Hi-Fidelity PCR Mastermix (Thermo Scientific).

Amplicons of the native Tp1 gene were digested with *Bam*HI and *Xho*I FastDigest restriction enzymes (Table 1) (Thermo Fisher Scientific) and directly ligated into either pYD1 or pYES2/NTC plasmids with compatible restriction ends using T4 DNA ligase (Thermo Fisher Scientific). The pYD1 expression vector is a low copy number plasmid designed for surface display of proteins by high-jacking the Aga1-Aga2 system of the yeast. Proteins are expressed in-frame linked to Aga2, with a 5’ V5 marker and a 3’ Xpress marker. In contrast, the pYES2/NTC expression vector is a high copy number plasmid designed for intracellular protein expression in yeast (Suppl. Table 1). Plasmids were transformed into chemically competent *E. coli* DH5α, DH10B (New England Biolabs) or TOP10 cells (Invitrogen), and transformed *E. coli* were plated. Subsequent selection was carried out on LB (Oxoid) supplemented with ampicillin (100 µg mL^−1^, Sigma Aldrich). Recombinant plasmids were prepared from positive *E. coli* clones and Sanger sequenced using plasmid specific primers for validation.

Amplicons of the native Tp2 gene were ligated blunt-end into pJET1.2 (Thermo Scientific) and transformed into chemically competent *E. coli* DH10B. Recombinants were sequenced and used as template for Tp2 amplification and ligation into pYD1 as described for Tp1 above. The native Tp9 gene was amplified from pcDNA3_Tp9 (kindly provided by Dr. N. MacHugh, Roslin Institute, Edinburgh, UK). Amplicons were cloned directly into pYD1 as described above. Yeast codon-optimised sequences of the Tp1 and Tp2 gene were synthesized by GeneArt (Thermo Fisher Scientific) and codon-optimised sequences for expression of Tp2 and Tp9 gene in eukaryotic cells were kindly provided by Dr. T. Connelley and Dr. N. MacHugh (Roslin Institute, Edinburgh, UK).

Eight EBY100 and INVSc1 yeast clones were selected and each cultured in 5 mL YNB-CAA broth and SC-U broth for 16-18h, respectively. Plasmids were extracted using the Qiagen Miniprep Kit using 425-600 μm glass beads (Sigma Aldrich) for mechanical lysing of yeast cells. Beads were added to the P1 solution and yeast cells were vortexed for 5 min, prior to the addition of P2 solution. Subsequent steps were conducted as described in the kit manual. Extracts were analysed for inserts by PCR with either pYD1_F/R primers, or pYES2/NTC_F/R primers (Table 1), and yeast clones containing the correct inserts were analysed subsequently for antigen expression.

### Analysis of Tp antigen expression by flow cytometry

*T. parva* antigen genes cloned into the pYD1 plasmid were expressed and protein displayed on the yeast cell surface was detected using an antibody to the C-terminal V5 epitope. Yeast expression was induced according to the manufacturer’s recommendations. Briefly, 18-20h yeast cultures from each of the eight EBY100 clones and one clone containing the empty pYD1 vector were prepared in separate 10 mL YNB-CAA 2% glucose at 30°C, 220 rpm to an OD_600_ of 2. Cells were pelleted, resuspended in induction medium YNB-CAA 2% galactose to an OD_600_ of 1 and grown for 48h at 20°C, 220 rpm. Cells equivalent to an OD_600_ of 0.5 were collected in duplicate at 0h, 24h, 48h and labelled with 4 ng μL^−1^ mouse V5 specific primary antibody (Invitrogen) followed by 4 ng μL^−1^ goat-anti-mouse-AF488 (Thermo Scientific). Cells were resuspended in 1 mL FACSFlow buffer (BD Biosciences) and analysed by flow cytometry using a FACSAria II (BD Biosciences) with seven PMTs with standard filters (LP: 520 nm, BP: 530/30 nm; LP: 556 nm, BP: 576/26 nm; LP: 595 nm, BP: 601/20 nm; LP: 655 nm, BP: 695/40 nm; LP: 735nm, BP: 780/60 nm; BP: 660/20 nm; LP: 735 nm, BP: 780/60 nm), a 488 nm Argon and a 633 nm Helium-Neon Laser. Acquisition settings were defined using unstained samples by adjusting voltage to the third logarithmic (log) decade of FSC and Doublet Discrimination from the FSC-Width versus SSC-width signal. FSC was used as trigger signal. Samples were acquired in the 530/30 nm fluorescence channel, using dot plot: SSC versus AF488 fluorescence. Events were recorded using BD FACSDiva software (Version 7, BD Biosciences), data were exported in FCS 2.0 file format and analysed using FlowJo (Version 10, Windows Operating Software) and expressed using two parameters: (a) percentage of cells AF488-positive cells and (b) Median Fluorescent Intensity (MFI).

Specificity controls used in flow cytometry experiments include unstained cells, isotype control antibody (mouse IgG2a, Thermo Fisher Scientific), and cells stained only with the secondary antibody. EBY100 cells were analysed at 0h, 24h and 48h, and included clones containing an empty pYD1 vector or clones expressing either Tp2 (native or yeast codon-optimised) or Tp9 (native or *Bos taurus* codon-optimised) at 0h, 24h and 48h.

### Quantification of antigens expressed in yeast

To quantify yeast surface displayed antigens, the two disulphide bonds between Aga1 and Aga2 were cleaved as described previously with modifications^33^. To do so, 10 ml of induced recombinant EBY100 cells were pelleted and frozen. Each pellet was resuspended in 2 mL dithiothreitol (DTT) extraction buffer (2 mM DTT, 25 mM Tris-HCl, pH 8) and incubated for 2h at 4°C on a rocking platform. Cells were separated by centrifugation at 6,000 x g for 5 min and cleaved proteins in the supernatant were concentrated and dialyzed with cleavage buffer using an Amicon filter of 10kD MWCO (Millipore) according to manufacturer’s instructions. Proteins were precipitated with the ProteoExtract Protein Precipitation Kit (Calbiochem) and quantified by Coomassie (Bradford) Protein Assay (Thermo Scientific). Equal amounts of protein for each set of samples (e.g. different time points of one clone) were loaded for SDS-PAGE and Western blot.

Antigens expressed in pYES2/NTC were detected either by a C terminal V5 or HisG epitope. Yeast expression was induced as described above and sampled at 0h, 4h, 8h, 12h or 14.5h, and 24h to test for antigen expression. Cells were lysed with YPER+dialyzable yeast protein extraction reagent (Thermo Scientific) and Halt Protease Inhibitor Cocktail (Thermo Scientific) according to the wet cell weight. Yield of protein was quantified using Coomassie (Bradford) Protein Assay (Thermo Scientific). The amount of protein for each clone at different time points was standardized, then precipitated with the ProteoExtract Protein Precipitation Kit (Calbiochem). Equal amounts of protein for each set of samples (e.g. different time points of one clone) were loaded for SDS-PAGE and Western blot.

### SDS-PAGE and Western Blot analysis

SDS-PAGE was carried out using precast 4-20% gradient polyacrylamide gels (Expedeon) and stained with InstantBlue (Expedeon). The PageRuler Prestained NIR Protein Ladder (Thermo Scientific) was included. Proteins were transferred to PVDF membranes (Millipore) in run blot buffer (Expedeon) and 20% methanol (VWR), washed in TBS-T buffer and blocked in TBS-T with 5% skim milk (Sigma). Blots were incubated overnight at 4°C with agitation in blocking buffer and anti-HisG-HRP (1:5000, Invitrogen), or anti-V5-HRP (1:5000, Invitrogen). Chemiluminescent detection of blots was conducted with Luminata Forte (Millipore) in a G-box illuminator with Epi Red and FRLP filters (Syngene) for visualizing ladder and samples, respectively.

### Inactivation and viability assay of yeast

For *in vitro* and *in vivo* experiments, yeast was used in a heat-inactivated and freeze-dried form. Yeast cultures were heat inactivated at a dilution of 1:20 in 1xPBS at 56°C for 1h. Inactivated cell pellets were freeze-dried in a Lyodry Compact Benchtop Freeze Dryer (MechaTech Systems) for 5-18h. Viability of cells was assayed by culturing a 10 μl loopful of freeze-dried cells in 10 mL YPD broth at 30°C, 250 rpm for three days. In case of growth, positive broth cultures were streaked onto YPD plates and incubated at 30°C for 3 days to ensure pure cultures were obtained.

### Immunisation of cows

Three castrated male Aberdeen Angus/Holstein Friesian cross calves (RVC, Bolton Park Farm) were used. Animals were dewormed and tested negative for Bovine Viral Diarrhoea Virus (BVDV), *Mycobacterium bovis* and Bovine Herpes Virus 1 (Infectious Bovine Rhinotracheitis, IBR). All calves were born within 18 days of each other and identified by the last three digits of their ear tag number: 224, 230, and 232. ITM was used for immunisation. Each calf, eight to ten weeks old, was inoculated with 250 μL of *T. parva* Muguga (TpM) sporozoite stabilate S71. Long lasting tetracycline was administered intramuscularly (i.m.) at the time of infection. The drug dose was adjusted to the weight of the animals according to the manufacturer’s recommendations. Four weeks after the first treatment each calf was inoculated with 3×10^6^ autologous TpM schizont infected bovine PBMCs to boost the immunity. Parasitized cells were suspended in 0.5 mL 1xPBS and applied subcutaneously (s.c.) in front and slightly above the right prescapular lymph node. No additional drug treatment was given. Animal treatment and sampling was approved after initial AWERB assessment and under Home Office licence (PPL7009059 and PPL60/4394)

### MHC class I genotyping

The protective cell-mediated immune response to *T. parva* is dominated by an MHC class I restricted response^34^. The range of expressed MHC class I allelic diversity was determined in experimental animals 224 and 232 as described previously^35^ with minor modifications^36^. Total RNA was extracted using the Qiagen RNeasy Mini kit from either cell lines of known genotypes as positive controls^37^ or lymphocytes of TpM infected calves. Contaminating DNA was removed with the Turbo DNA-free™ kit (Applied Biosystems) before cDNA was synthesized with the AMV LongAmp^®^ Taq RT-PCR kit (New England Biolabs). A 500bp fragment covering the second and third exons common to all polymorphic MHC class I genes as amplified from each cDNA using NEB Go Taq polymerase and ovine MHC class I generic primers 416 and Cr as described in^36^. Each fragment was gel purified and sequenced in both directions to ensure amplification of a range of MHC class I alleles. An optimized number of 24 cycles was used to eliminate amplification of chimeric alleles. Amplified fragments purified by gel electrophoresis, were cloned in the Promega T-easy vector. Successfully transformed *E. coli* were selected on ampicillin agar plates (100 μg mL^−1^). Forty bacterial colonies from each plate were screened by PCR for the presence of inserts. Subsequently, 38 clones from animal 224 and 36 clones from animal 232 were Sanger sequenced in both directions. Individual MHC class I sequences were assembled from overlapping forward and reverse sequences of each clone using the SeqMan Pro Programme within the DNAstar Lasergene11 package. For validation, individual MHC class I sequences derived from a minimum of two independent clones were BLAST searched against the NCBI and IPD-MHC databases (https://www.ebi.ac.uk/ipd/mhc/). The nucleotide sequence of a novel MHC class I transcript identified in animal 232 was submitted to the European Nucleotide Archive (ENA) and received the following database accession, (LR994465).

### Generation of *T. parva* Muguga infected cell lines

Parasitized cell lines were generated by *in vitro* infection of bovine PBMCs obtained from the calves before ITM immunisation^38^. Cell lines were maintained at 37°C under 5% CO2 in RPMI-1640 Glutamax cell culture medium (Life Technologies, Gibco, Paisley, UK) supplemented with 10% FCS (GIBCO), penicillin-streptomycin (100 units mL^−1^ and 100 μg mL^−1^, final concentrations respectively, Sigma-Aldrich, Dorset, UK) and 50 μM (final concentration) of 2-mercaptoethanol (Sigma Aldrich).

### Stimulation and enrichment of CD8^+^ T cells

Antigen specific CD8^+^ effector T cells were expanded from whole blood after ITM immunisation. Expansion involved stimulation with gamma irradiated (100 Gy) autologous TpM lines. Effector and stimulator cells were co-cultured at an effector to stimulator ratio of 20:1 in 2 mL complete culture medium (5×10^6^ effector: 2.5×10^5^ stimulator cells per well of a 24 well plate). Co-cultures were incubated at 37°C in 5% CO2 for one week, harvested and purified over a Ficoll-Paque density gradient (GE Life Sciences). Viable effector cells were re-stimulated at an effector to stimulator ratio of 10:1 (2.5×10^6^ effector: 2.5×10^5^ stimulator cells) for another week and enriched for CD8 by complement mediated lysis of CD4^+^ and γδ-T cells as described^36^. Ficoll-Paque purified cells were plated at a concentration of 5×10^3^ cells per well in a 96 well plate and re-stimulated with 1×10^3^ irradiated TpM cells in the presence of 100 units mL^−1^ of recombinant human IL-2 (Proleukin®, Novartis) for future use.

Successful enrichment of CD8^+^ or CD4^+^ T cells was confirmed by flow cytometry. The effector cell suspension was stained with 4 μg mL^−1^ of primary antibodies (Table 2) targeting four different T cell populations as well as 4 μg mL^−1^ of goat-anti-mouse IgG AF488 secondary antibody (Thermo Scientific). Acquisition and data analysis were performed as mentioned above.

**Table 2.** Summary of primary antibodies used for confirmation of antigen surface expression

| Reactivity | Antibody (source) | Final concentration<br>($\mu\text{g mL}^{-1}$) |
| --- | --- | --- |
| Mouse Anti-CD8, bovine | IL-A105 (Sigma-Aldrich, 91072535) | 4 |
| Mouse Anti-CD4, bovine | IL-A11 (Sigma-Aldrich, 91072511) | 4 |
| Mouse Anti-WC1, bovine | Clone CC15 (Invitrogen, MA516616) | 4 |
| Mouse Anti-CD335, bovine | NKp46 (Biorad, AKS1) | 4 |

### Cytotoxicity assay

The CytoTox96^®^ Non-Radioactive Cytotoxicity Assay (Promega, Cat no. G1780) was performed according to the manufacturer’s instructions (G1780 Literature # TB163). Briefly, duplicate wells were seeded with TpM infected target cells at a concentration of 1×10^4^ cells per well. Enriched CD8^+^ T effector cells were added in decreasing concentrations to achieve effector to target ratios from 20:1 to less than 1:1 per well. Co-cultures were maintained in phenol red-free RPMI-1640 medium (GIBCO) with 2% FCS and incubated at 37°C in 5% CO_2_. The percentage of cytotoxicity was calculated as recommended by the protocol using Excel (software version 2019 for Windows10). Briefly, values were first corrected for culture medium background. The experimental LDH release (OD_490_) was divided by the maximum LDH release (OD_490_) and further multiplied by 100.

### Assessment of cytokine production and cytokine producing cells

IFNγ production was measured to assess the response of bovine PBMCs or enriched CD8^+^ T cells to yeast expressing Tp antigens. PBMCs were either freshly isolated from whole blood or from cells archived under liquid nitrogen. Enriched CD8^+^ T cells were generated as described above and used on days 10 to 17 post stimulation. Autologous PBMCs were thawed from Nitrogen stocks, while TpM infected cells were used as a positive control. Effector (2.5×10^4^) and yeast (2.5×10^4^) cells were co-cultured in 96-well plates at 37°C in 5% CO_2_ for 72h. PBMC (5×10^3^) were added where enriched CD8^+^ T cells represented the effector cells. Cell culture supernatants were analysed with the bovine IFNγ ELISA development kit (MABTECH, Cat no. 3119-1H-20) according to the manufacturer’s instructions. Optical density was measured in a SpectraMax M2 plate reader (Molecular Devices).

Ethanol-activated low fluorescent PVDF 96-well plates (Millipore) were coated with monoclonal antibodies to bovine IFNγ and IL-2 (MABTECH). Duplicate wells were seeded with 1×10^5^ cells per well of cryopreserved bovine PBMCs (rested overnight before use) that contained either complete culture medium (RPMI-1640 + 10% FBS, GIBCO), yeast cells expressing either empty plasmid or Theileria antigens at 2×10^5^ cells per well, or TpM infected cells at 1×10^4^ cells per well. Positive control wells containing PMA (50 ng mL^−1^) and Ionomycin (1 μg mL^−1^) (Sigma-Aldrich) were seeded with a reduced number of PBMCs, at 2.5×10^4^ cells per well. Additionally, wells containing only TpM infected cells were seeded. Plates were incubated at 37°C in 5% CO_2_ for 46h. Subsequently, plates were washed with 1xPBS and incubated with detection antibodies, IFNγ-BAM and IL-2-biotin (MABTECH). This was followed by incubation with fluorophore labelled secondary antibodies, anti-BAM-490 and streptavidin-550 (MABTECH). Finally, plates were incubated with a fluorescence enhancer (MABTECH). Cytokine producing cells were counted using an automated fluorescence plate reader (AID-iSpot reader, Autoimmun Diagnostika GmbH). Antigen-specific responses were calculated by deducting the number of spots formed in the background wells from the spots developed in response to the yeast, Theileria antigens and TpM infected cells. For the wells incubated with TpM infected cells the number of spots from TpM background wells were also deducted.

### Bacterial expression and purification of recombinant *T. parva* schizont antigen Tp2

Expression of full-length native Tp2 cloned into pET-19b was initially tested in *E. coli* BL21 Star (DE3) (Thermo Scientific) and SHuffle (New England BioLabs). For expression screening, positive transformants of both strains were grown in 10 mL LB medium (1% tryptone, 0.5% yeast extract, 0.5% NaCl; Sigma Aldrich) supplemented with ampicillin (100 μg mL^−1^, Melford Labs) for 3h at 37°C and induced with IPTG for 14h overnight at 20°C. Cells were harvested by centrifugation at 4,000 x g for 15 min at 4°C and resuspended in 2 mL of purification buffer (20 mM sodium phosphate + 0.5 M NaCl, pH 7.5). The cell suspension was sonicated for 2 min (10s on, 10s off, 30% amplitude) to collect the total cell protein fraction (T). Another centrifugation at 4,000 x g for 30 min was followed by collection of the supernatant containing the soluble protein fraction (S). Expression samples were checked on SDS-PAGE 4-20% gradient gels (NuPage) stained with InstantBlue (Expedeon).

Large scale expression was performed as described previously^39,40^. Essentially, Tp2 was expressed in *E. coli* BL21 Star (DE3) (Thermo Scientific) cells using overnight induction at 20°C. Soluble protein, after cell lysis with a French Press, was purified using enterokinase digest, HisTrap HP, and 5 mL affinity columns (GE Healthcare) on an AktaPure chromatography system. Purified Tp2 in the eluted fractions was identified by SDS-PAGE and Western blotting as described above.

Fractions containing Tp2 were pooled and concentrated to 5 mL for clean up by preparative Size Exclusion Chromatography (SEC) (Generon ProteoSEC 3-70 kD) (SEC buffer: 20 mM PO4, 250 mM NaCl, 0.02% NaN3, pH 7.5). Monomer and dimer species were pooled separately and concentrated to 2.5 mL using VivaSpin 20 centrifuge concentrators with 5 kD MWCO by centrifugation at 2,000 x g. Samples were buffer exchanged into NMR buffer (20 mM sodium phosphate pH 7.0, 50 mM NaCl, 0.02% NaN3) using PD10 desalting columns (GE Healthcare). Samples were concentrated further to a volume of ~300 μL, mixed with 5% of D2O (Sigma Aldrich), adjusted to a pH of 7.25 with HCl and transferred into a 5 mm NMR tube (Shigemi). Protein for a ^15^N labelled sample was expressed in modified minimal media as described previously^41,42^ with purification and sample preparation as described for the unlabelled samples. NMR spectra were recorded on Bruker Avance III 700 and 800 MHz spectrometers equipped with cryoprobes. 1D and 2D ^1^H-^15^N HSQC spectra were recorded with standard pulse-sequences provided by the manufacturer.

Molecular weights of apparent monomer and dimer fractions were characterized by Size Exclusion Chromatography-Multiangle Light Scattering (SEC-MALS) using a FPLC system (Shimadzu) with integrated degassing chamber as well as separate scattering and refractive index detectors (Wyatt technologies). Data analysis was conducted using the manufacturer’s software. Circular Dichroism (CD) analysis of both the monomer and dimer fraction was performed with an Applied Photophysics Chirascan Plus instrument. Far UV CD spectra were recorded using a rectangular 0.5 mm Quartz Suprasil cuvette (Starna Scientific Ltd.) with a bandwidth of 1 nm, 1 nm stepsize and 1.5 s accumulation time per point. Spectra for each sample were recorded at room-temperature for the far UV (200-250 nm) and near UV (250-400 nm) followed by a Melting Curve monitored in the far UV region. Samples were measured in NMR buffer with and without DTT.

### Immunogenicity of *S. cerevisiae* expressing Tp2

The immunogenicity of Tp2 stably expressed on the surface of *S. cerevisiae* was assessed by measuring the antibody response in mice. Two groups of ten female, five-week-old BALB/c mice were immunised orally with 30 mg of inactivated, freeze-dried yeast dissolved in 100 μL of 1xPBS and containing either the empty plasmid or extracellularly expressing Tp2 on day 0, 15, 30, 45 and 60. Blood was collected for serum preparation on day (−4) and at the end of the trial on day 90. Blood was stored at −20°C until further use. This study was conducted by Clinvet (Bloemfontein, RSA) and approved by the ClinVet Institutional Animal Care and Use Committee (IACUC; study number CG411-CV18/178).

### Measurement of total IgG and Tp2 specific IgG in mouse serum

The total IgG concentration in mouse serum was determined using the IgG (Total) Mouse Uncoated ELISA Kit (Invitrogen, Cat no. 88-50400) according to the manufacturer’s instructions. Serum samples were diluted 1:10,000 before addition to ELISA plates. The optical density was measured at 450 nm with a Multiscan FC Plate Reader (Thermo Scientific).

To assess Tp2 specific IgG production, Nunc MaxiSorp™ Flat bottom 96-well ELISA plates were coated with Tp2 monomer (10 μg mL^−1^, 100 μL per well) at 4°C overnight. After blocking (1xPBS + 0.1% Tween™20 + 1% BSA) for 2h at RT, mouse serum samples (diluted 1:30) were added to each well and incubated with cross-absorbed anti-mouse IgG HRP polyclonal antibody (diluted 1:10,000) (STAR117P, Bio-Rad) for 2h at RT on a Multiscan FC Plate Reader (Thermo Scientific) at medium shaking speed. After four washes in 1xPBS with 0.05% Tween™20, plates were developed with TMB substrate solution (Invitrogen) for 20 min at RT. The enzymatic reaction was stopped by adding 2N H_2_SO_4_ (R&D Systems) to each well and the plates read at 450 nm using a Multiscan FC Plate Reader (Thermo Scientific). ELISA results were presented as normalised OD_450_ with the mean OD_450_ values of blank wells subtracted from the sample well values.

### Statistical analysis

Data from fluorospot assay, APC and yeast cell titration, total IgG serum ELISA and Tp2 specific IgG serum ELISA was analysed using GraphPad Prism (v 8.1.1) (GraphPad Software, San Diego). For the PBMC fluorospot assay, the corrected number of spots for the pYD1 control and pYD1_Tp2 were compared by two independent samples t-test. For APC and yeast cell titration, data was transformed to log scale for subsequent non-linear regression analysis. For analysis of murine IgG ELISA data, the absorbance values of sample wells were corrected for unspecific signal from blank wells. For total serum IgG analysis, the absorbance values of the standard curve were plotted and the concentration of total IgG in each sample interpolated in GraphPad Prism. The increase in total serum IgG over time was calculated for both groups and a two way independent samples t-test performed. For murine Tp2 specific serum IgG analysis, the increase in absorbance over the study period was calculated for both groups and a two way independent samples t-test performed to compare both means.

## Results

### Expression of Tp1, Tp2 and Tp9 in yeast

Five *T. parva* schizont expressed genes coding for Tp1, Tp2, Tp9, Tp10 and N36 antigens were initially selected for expression in yeast. Each gene was cloned into the pYD1 vector and the level of surface antigen expression determined by flow cytometry. Screening of eight yeast clones per gene at 24h and 48h identified clone specific variation in surface expression (Fig. 1). Clones with the highest surface expression were selected for subsequent experiments (Fig. 1/ Table 3). Codon optimisation had no impact on expression levels, hence native sequences for each protein were subsequently used.

**Figure 1:**
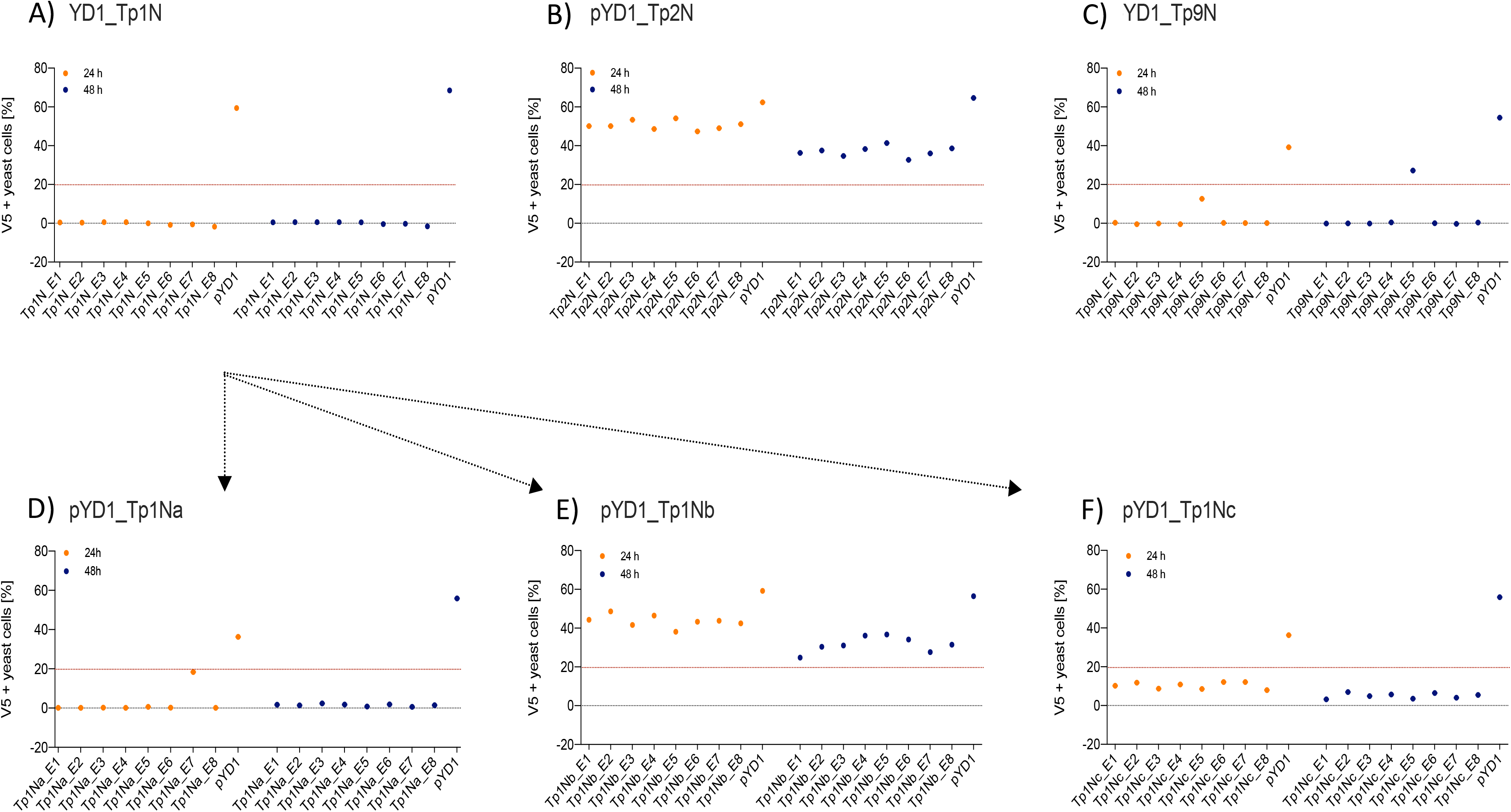
Positive N-terminal V5 surface staining of EBY100 yeast expressing different *T. parva* schizont antigens of native sequence (A) Tp1, (B) Tp2, and (C) Tp9. Eight clones were analysed by flow cytometry after 24h (orange) and 48h (blue) post induction for each antigen. Results displayed represent one of two independent experiments and are corrected for positive staining of EBY100 yeast at 0h post induction. Red lines were included to alleviate the comparison of antigen expression. Unsuccessful identification of native Tp1 was followed by analysis of three Tp1 fragments (D) Tp1Na, (E) Tp1Nb, and (F) Tp1Nc. Clones stably expressing comparably high amounts of antigen could then be selected for all antigens.

**Table 3.**
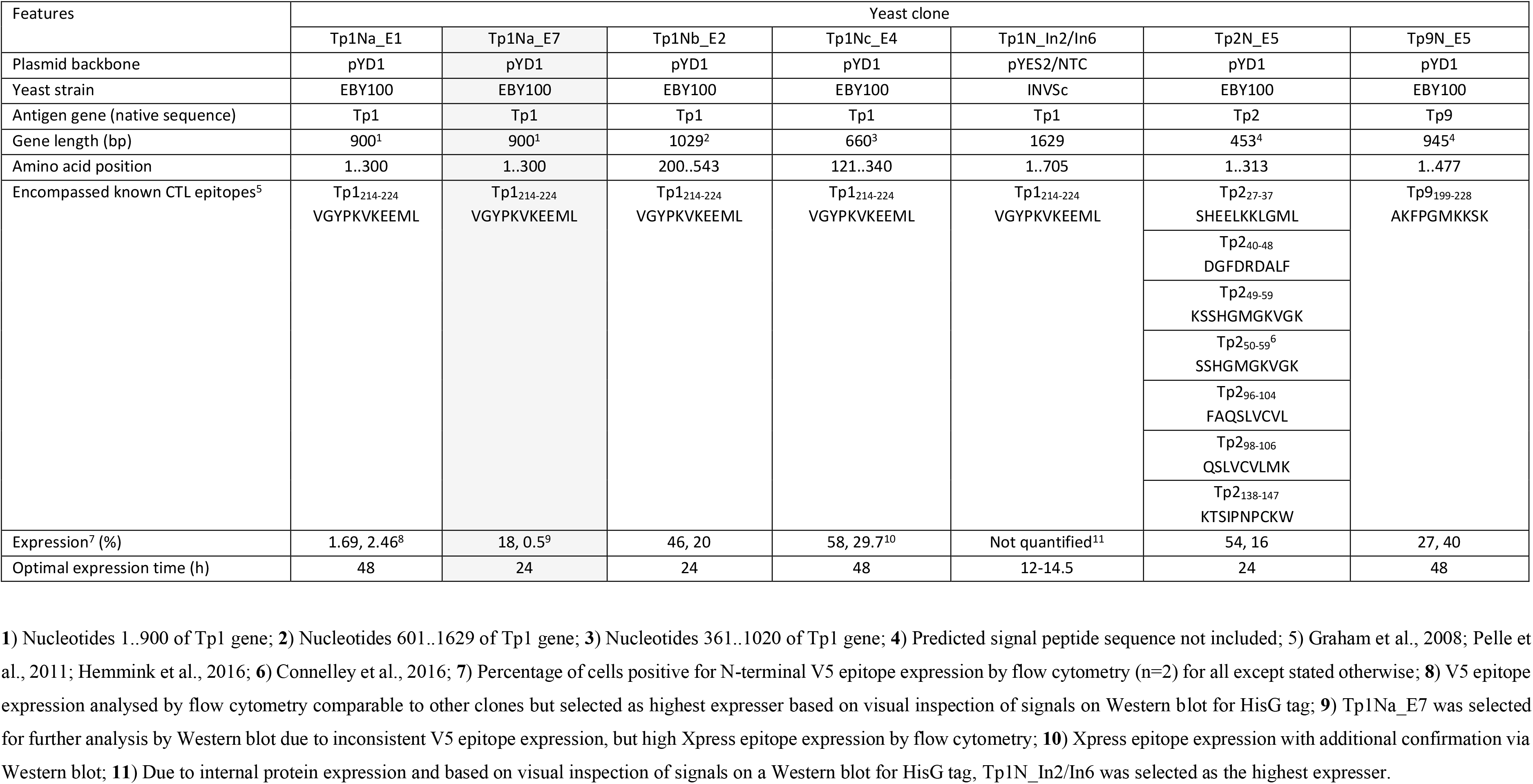
Summary of yeast clones expressing T. parva Muguga antigens

| Features | Yeast clone |  |  |  |  |  |  |
| --- | --- | --- | --- | --- | --- | --- | --- |
|  | Tp1Na_E1 | Tp1Na_E7 | Tp1Nb_E2 | Tp1Nc_E4 | Tp1N_In2/In6 | Tp2N_E5 | Tp9N_E5 |
| Plasmid backbone | pYD1 | pYD1 | pYD1 | pYD1 | pYES2/NTC | pYD1 | pYD1 |
| Yeast strain | EBY100 | EBY100 | EBY100 | EBY100 | INVSc | EBY100 | EBY100 |
| Antigen gene (native sequence) | Tp1 | Tp1 | Tp1 | Tp1 | Tp1 | Tp2 | Tp9 |
| Gene length (bp) | 900<sup>1</sup> | 900<sup>1</sup> | 1029<sup>2</sup> | 660<sup>3</sup> | 1629 | 453<sup>4</sup> | 945<sup>4</sup> |
| Amino acid position | 1..300 | 1..300 | 200..543 | 121..340 | 1..705 | 1..313 | 1..477 |
| Encompassed known CTL epitopes<sup>5</sup> | Tp1<sub>214-224</sub><br>VGYPKVKEEML | Tp1<sub>214-224</sub><br>VGYPKVKEEML | Tp1<sub>214-224</sub><br>VGYPKVKEEML | Tp1<sub>214-224</sub><br>VGYPKVKEEML | Tp1<sub>214-224</sub><br>VGYPKVKEEML | Tp2<sub>27-37</sub><br>SHEELKKLGML | Tp9<sub>199-228</sub><br>AKFPGMKKSK |
|  |  |  |  |  |  | Tp2<sub>40-48</sub><br>DGFDRDALF |  |
|  |  |  |  |  |  | Tp2<sub>49-59</sub><br>KSSHGMGKVGK |  |
|  |  |  |  |  |  | Tp2<sub>50-59</sub><sup>6</sup><br>SSHGMGKVGK |  |
|  |  |  |  |  |  | Tp2<sub>96-104</sub><br>FAQSLVCVL |  |
|  |  |  |  |  |  | Tp2<sub>98-106</sub><br>QSLVCVLMK |  |
|  |  |  |  |  |  | Tp2<sub>138-147</sub><br>KTSIPNPCKW |  |
| Expression<sup>7</sup> (%) | 1.69, 2.46<sup>8</sup> | 18, 0.5<sup>9</sup> | 46, 20 | 58, 29.7<sup>10</sup> | Not quantified<sup>11</sup> | 54, 16 | 27, 40 |
| Optimal expression time (h) | 48 | 24 | 24 | 48 | 12-14.5 | 24 | 48 |

**1)** Nucleotides 1..900 of Tp1 gene; **2)** Nucleotides 601..1629 of Tp1 gene; **3)** Nucleotides 361..1020 of Tp1 gene; **4)** Predicted signal peptide sequence not included; **5)** Graham et al., 2008; Pelle et al., 2011; Hemmink et al., 2016; **6)** Connelley et al., 2016; **7)** Percentage of cells positive for N-terminal V5 epitope expression by flow cytometry (n=2) for all except stated otherwise; **8)** V5 epitope expression analysed by flow cytometry comparable to other clones but selected as highest expresser based on visual inspection of signals on Western blot for HisG tag; **9)** Tp1Na\_E7 was selected for further analysis by Western blot due to inconsistent V5 epitope expression, but high Xpress epitope expression by flow cytometry; **10)** Xpress epitope expression with additional confirmation via Western blot; **11)** Due to internal protein expression and based on visual inspection of signals on a Western blot for HisG tag, Tp1N\_In2/In6 was selected as the highest expresser.

Consistently very low levels of expression were detected for Tp1 while identification of surface expression of Tp10 and N36 failed. As Tp1, Tp10 and N36 genes are considerably larger (1,629bp, 1,329bp and 1,662bp, respectively) compared to Tp2 (453bp) and Tp9 (945bp), two strategies were developed in attempts to increase expression of Tp1, whereas expression of Tp10 and N36 was abandoned. The first strategy involved the expression of Tp1 subunits Tp1Na, Tp1Nb and Tp1Nc, in pYD1 in a manner that preserved the antigenic epitope in all fragments (Suppl. Fig. 1). Resulting clones showed low and variable V5 (0.5-18%) as well as Xpress expression (3-9%) (Fig. 1D-F). Tp1Na_E7 was selected as the highest expressing clone. In contrast, V5 expression of Tp1Nc was low (3-12%; Fig. 1F), but Xpress expression relatively high (48-63%; Table 3). Clone Tp1Nc_E4 was subsequently selected (Fig. 1F/ Table 3). All Tp1Nb clones showed similar V5 expression (25-49%, Fig. 1E/ Table 3) with Tp1Nb_E2 displaying stable and moderate V5 expression similar to Tp2N_E5 and Tp9N_E5 clones (Fig. 1B and C/ Table 3).

The low detection of Tp1Na could have been a result of inaccessible epitopes within the surface displayed Tp1Na and therefore poor detection. Consequently, attempts were made to cleave surface displayed Tp1Na at 24h and 48h after induction of the yeast cell surface. Western blot analysis using an anti-V5-antibody labelled with HRP only resulted in bands of 50kD, 55kD and 80kD in clone Tp1Na_E1 after 48h induction (Fig. 2A). Protein cleaved from the Tp1Nc_E4 clone was included as a positive control. Here, a band at 90kD could be observed (Fig. 2B). Theoretical molecular weights of Tp1Na (50kD) and Tp1Nc (41kD) suggest that the proteins could be either glycosylated or aggregated.

**Figure 2:**
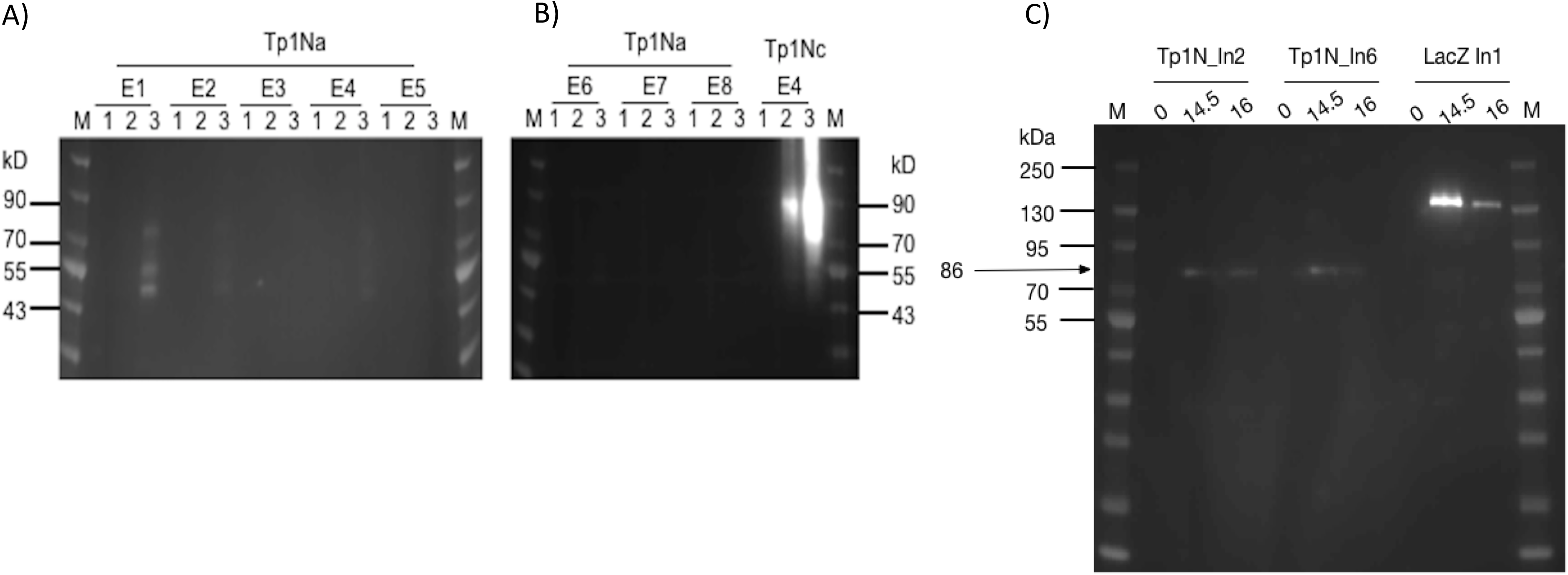
A) and B) Western blot analysis of yeast surface displayed Tp1Na and Tp1Nc recombinant proteins. DTT-cleaved proteins from eight clones of Tp1Na and one clone of Tp1Nc were adjusted in concentration so that each set of samples at 0h, 24h and 48h (indicated by 1,2,3 respectively) were standardised to the same amount. AntiAnti-HisG-HRP antibody was used. M: PageRuler pre-stained NIR Protein Ladder from Thermo Scientific. Image generated by merging images from exposure to near infra-red and no light. Clone Tp1Na_E1 and Tp1Nc_E4 were selected for further experiments. C) Western blot of Tp1N_In2, Tp1N_In6 and LacZ_In1 yeast clones, were LacZ_In1 is a positive control. Cultures were induced for 14.5h and 16h and lysed. Proteins were purified, precipitated and standardised to 20 mg per well. Recombinant protein was detected using anti-HisG-HRP antibody (1:2,500).

For the second strategy, the full-length Tp1N gene was ligated into pYES2/NTC and introduced into EBY100. Intracellular Tp1N expression of eight clones was assessed by anti-HisG Western blot analysis at five different time points post induction. Clones Tp1N_In2 and Tp1N_In6 showed equally high and reproducible expression at 14.5-16h with a major band at 86kD band (Fig. 2C). The theoretical molecular weight of recombinant Tp1N is 69kD.

### MHC class I genotyping of cows reveals differences in MHC class I haplotypes

Previous experiments have highlighted variation in the CD8^+^ cytotoxic T cell (CTL) response of cows to different Tp antigens^43,44^. The possibility of immunodominance associated with MHC class I restricted CD8^+^ cytotoxic T cell responses was evaluated by genotyping animals 224 and 232 at functional MHC class I loci. A broad range of alleles was detected at four of the six predicted MHC class I loci 1, 2, 3 and 4 (Table 4) in cow 224. Three of these alleles have previously been associated with serologically defined MHC class I specificities A14 and/or A15^45^ which are known to restrict the CTL response to Tp9^46^. In contrast, the distribution of MHC class I alleles derived from animal 232 was more limited. Four sequences appear either identical or are very similar to alleles assigned to locus 3. A novel allele was identified in animal 232 (Accession LR994465) along with two non-classical MHC class I genes. Three of the classical alleles identified in animal 232 are associated with known serological MHC class I specificities, but only one of these specificities was previously associated with CD8^+^ cytotoxic T cell responses to Tp9^46^.

**Table 4.** MHC class I alleles and associated serological specificities found in cows 224 and 232

| Cow | Allele | Differences | Number of sequenced clones with the allele | Associated serological specificity (SSP)* | Breed the allele is mostly detected in<sup>1</sup> |
| --- | --- | --- | --- | --- | --- |
| **224** | *BoLA-class I-1\*00901* | - | 7 | A15 | Holstein |
|  | *BoLA-class I-2\*02501* | - | 2 | A14/A15 | Holstein |
|  | *BoLA-class I-2\*03202* | - | 7 | - | Angus cross |
|  | *BoLA-class I-3\*06801* | - | 11 | - | Angus cross |
|  | *BoLA-class I-4\*02401* | - | 10 | A14/A15 | Holstein |
|  | *BoLA-class I-NC1\*0101* | 3 bp differences | 1 | - | - |
| **232** | *BoLA-class I-3\*00401* | 1 bp difference | 2 | A33 | Hereford |
|  |  | 100% identity to *Bibi-N\*01201* and<br>*BoLA Accn no L02832* |  |  |  |
|  | *BoLA-class I-3\*01001* | - | 5 | A10 | Boran |
|  | *BoLA-class I-3\*02701* | - | 8 | A20 | Holstein |
|  | *BoLA-class I-3\*03601* | - | 13 | - | Holstein |
|  | *BoLA-class I-NC2\*00103* | - | 2 | - | - |
|  | *BoLA-class I-NC3\*0101* | 2 bp differences | 3 | - | - |
|  | *BoLA-class I-New* | 31 bp differences<br>from *BoLA-2\*01601* | 2 | - | - |

\* Hammond et al. 2012

### *T. parva* antigens expressed in *S. cerevisiae* elicit an IFNγ response but not an IL-2 response from bovine PBMCs *in vitro*

Successful control of ECF involves induction of a MHC class I restricted CTL response that targets parasite-infected white blood cells. Here we evaluate the response to *T. parva* antigens stably expressed on the surface of yeast. PBMCs from three cows previously immunized by the ITM were incubated with yeast cells. The number of IFNγ and IL-2 producing cells was measured by fluorospot. The IFNγ response differed between the three animals (Fig. 3A-C). IFNγ producing cells were identified in all wells while IL-2 production was only observed in the positive control (Fig. 3D, IX). PBMCs from all three cows responded to yeast expressing Tp2 compared to yeast containing an empty pYD1 plasmid (green bars). This response was statistically significant for cow 224 (p<0.05). Furthermore, elevated numbers of IFNγ producing cells were identified in PBMC cultured with yeast expressing Tp1_Nc for cow 224 and 230, and for Tp9 for animal 232. However, none of these responses reached significance at p < 0.05. In contrast, yeast expressing the Tp1_Nb fragment had a suppressive effect with a decrease in IFNγ producing cells below the empty plasmid background in cows 224 and 232. To determine if the variation in response of PBMC from different cattle to Tp antigens is evidence for immune dominance associated with MHC class I restricted cytotoxic T cell response we then assessed whether Tp antigens expressed on *S. cerevisiae* were able to induce a CTL response.

**Figure 3:**
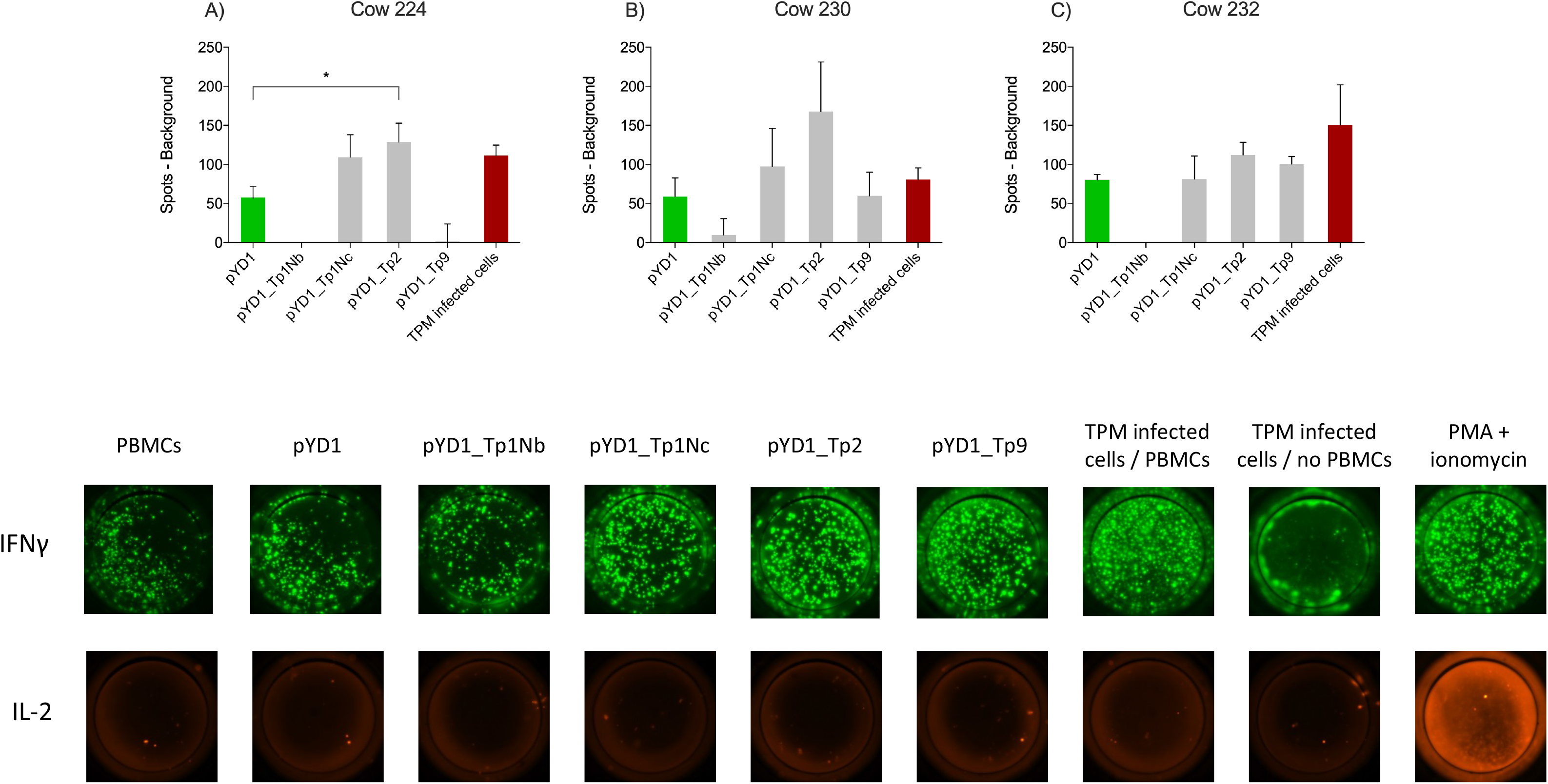
IFNγ responses by bovine PBMCs to yeast cells expressing *T. parva* schizont antigens determined by fluorospot analysis. PBMCs from cow A) 224, B) 230 and C) 232 were assayed in duplicate on four separate occasions. Data are shown as the mean number of spots (±SEM) generated in response to antigen minus the number of background spots generated by PBMCs alone. For the wells incubated with TpM infected cells the number of spots from TpM background wells was also deducted. Where the mean number of spots produced in response to Theileria antigens was less than background the number of spots was reported as zero. D) Typical IFNγ and IL-2 dual fluorospot images from bovine PBMCs incubated with EBY100 yeast cells expressing different *T. parva* schizont antigens. The empty plasmid pYD1 represented the negative control while TpM infected cells incubated with PBMCs as well as stimulation with PMA and ionomycin served as positive control. While an increased IFNγ response towards pYD1_Tp1Nc, pYD1_Tp2 and pYD1_Tp9 was observed, no IL-2 production was detected.

### *T. parva* antigens presented by *S. cerevisiae* induce MHC class I restricted CD8^+^ CTL responses *in vitro*

To verify this hypothesis, PBMC isolated from all three immunised cattle were stimulated with autologous, gamma irradiated TpM infected cell lines and enriched for CD8 or CD4 T cell markers. The frequency of cells expressing CD4, CD8, WC1 (γδ T cells) and NKp46 (NK cells) markers was assessed by flow cytometry which confirmed CD8^+^ or CD4^+^ T cell enrichment to 90% and 95%, respectively (Suppl. Fig. 2A). The cytotoxic activity of both enriched cell populations was assessed by quantitative measurements of lactate dehydrogenase (LDH) released upon cell lysis. Increasing concentrations of CD8^+^ T cells led to an increased lysis of TpM infected target cells. In contrast, CD4^+^ T cells showed almost no cytotoxic activity (Suppl. Fig. 2B and C). The cytotoxic activity of CD8^+^ T cells was restricted to autologous TpM infected cell lines. CD8^+^ T cells from cattle 224 effectively killed autologous target cells while heterologous TpM from animal 232 were not lysed (Suppl. Fig. 2D). Equally, effector cells from animal 232 only reacted to autologous target cells (Suppl. Fig. 2E). This could have occurred due to a MHC class I restricted T cell response in cattle with varying MHC class I haplotypes ^47^.

For subsequent assays, IFNγ produced by enriched, cytotoxic CD8^+^ T cells towards yeast expressing *T. parva* antigens was assessed as a marker for cell-mediated killing, similar as described earlier ^36^.

Initial experiments highlighted a strong IFNγ production by APCs causing a misleading background signal. Thus, both APCs (Suppl. Fig, 2F) and yeast cells (Suppl. Fig. 2G) were titrated to identify suitable numbers per well and optimise the signal-to-noise ratio.

From the range of *T. parva* antigens expressed in yeast, IFNγ production by CD8^+^ effector T cells was observed towards native Tp2 and Tp9 (Fig. 4). Successful stimulation only occurred in co-cultures of yeast, APCs and T cells indicating that the antigens had to be processed and presented to T cells by APCs. The magnitude and antigen specificity of the immune response varied between animals. While Tp9 induced a strong reaction from animal 224 (Fig. 4A), only a marginal response was shown from cow 232. In contrast, Tp2 elicited a marginal response from cow 224 but no detectable reaction from animal 232 (Fig. 4B). Both cows showed an increased IFNγ production towards TpM infected cells. The response to Tp9 by one of the cows supports the concept of using yeast as a vector to deliver Theileria antigens, however, the variability between animals and absence of response to many of the antigens suggests that the system requires further optimisation.

**Figure 4:**
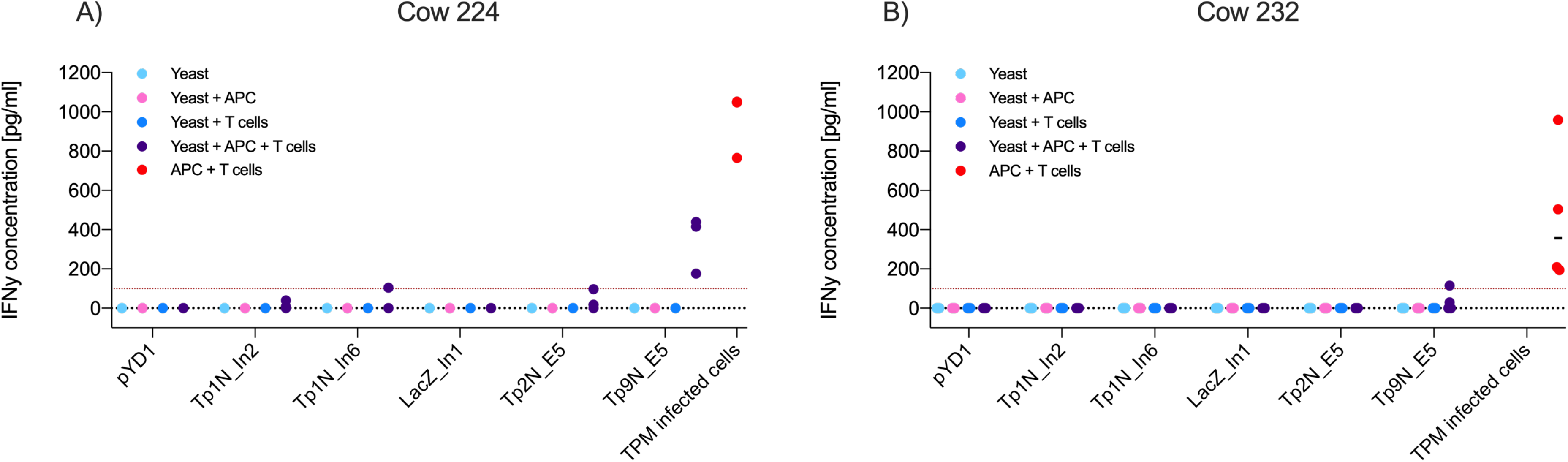
IFNγ production of antigen specific CD8^+^ effector T cells towards yeast expressing different *T. parva* antigens or towards TpM infected cells as positive control. Results displayed represent three/four independent experiments after correction for background signal. Red lines highlight an IFNγ concentration of 100 pg mL^−1^. Increased IFNγ production only occurred in co-cultures of yeast, CD8^+^ T cells and APCs, indicating that phagocytosis and processing of yeast by APCs was essential for T cell stimulation. A) While cow 224 responded to Tp9N and marginally to Tp1N_In6 as well as Tp2N, B) cow 232 only recognised Tp9N.

### Bacterially expressed Tp2 exhibits a stable and well-defined protein structure

Native Tp2 was the antigen showing consistently high extracellular expression in yeast, induction of a significant IFNγ response from bovine PBMCs during fluorospot analyses and stimulation of cytotoxic CD8+ T cells from one out of two animals. Previous identification of seven different CTL epitopes further supports Tp2 as an attractive candidate for evaluation of its immunogenic potential *in vivo* (Table 3). This first required the synthesis of this protein. Bacterial expression was initially tested in standard BL21 Star as well as SHuffle E. coli cells ^45^. The latter were engineered to produce soluble, folded proteins with correct disulphide bonds as Tp2 is predicted to contain six disulphide bonds based on the presence of 12 cysteines in its sequence ^48^. Interestingly, SDS-PAGE analysis of both total cell protein (T) and soluble protein (S) showed a distinct band at 22 kD in BL21 Star but not in SHuffle cells (Fig. 5A). The expected molecular weight of recombinant Tp2 was about 19.7 kD indicating the detected protein could be Tp2 which seems to be expressed mostly soluble in BL21 Star (Fig. 5A). Subsequent digestion of soluble Tp2 protein by enterokinase for 4h and 16h resulted in a band of 15-16 kD, which was very close to the expected molecular weight (Fig. 5B). Preparative expression of Tp2 was performed in BL21 Star in LB and M9 media producing about 10 mg L^−1^ LB and about 4 mg L^−1^ M9 of pure protein. Further purification by SEC following affinity chromatography revealed the protein in most fractions starting from the exclusion volume corresponding to an apparent molecular weight of >70 kD all the way to the approximate monomer molecular weight (uncut) of around 20 kD (Suppl. Fig. 3A). Peaks in the chromatogram at 70, 105 and 126 min correspond roughly to molecular weights of 68, 37 and 18 kD which are a good approximation of tetramer, dimer and monomer. To get a better MW estimate, the apparent dimer and monomer fractions were analysed by SEC-MALS (Suppl. Figure 3B). The apparent dimer eluted after 4.7 min while the scattering signal revealed a molecular weight of 43.9 +- 1.3 kD for the main peak. A shoulder at 5.5 min had a very noisy scattering signal which nevertheless was clearly identifiable as monomer. The apparent monomer eluted after 5.5 min and had a clear scattering signal of a monomer with a molecular weight of 23.8 +- 4.5 kD. No other species were detected in the monomer fraction.

**Figure 5:**
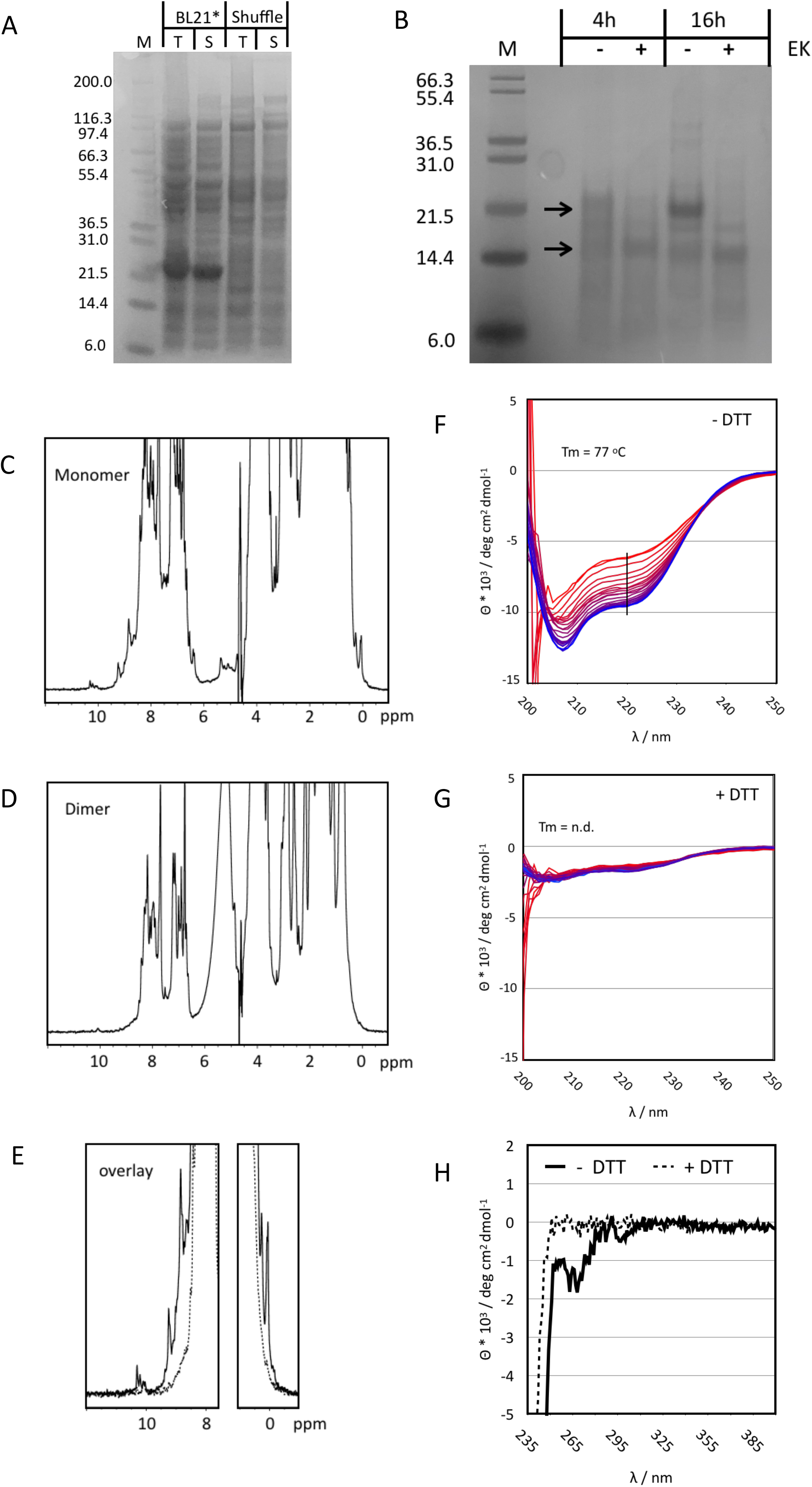
A) SDS-PAGE of small-scale test expression of Tp2 in BL21 STAR and SHuffle cells. Whole cell lysate is shown prior to (T) and after centrifugation (S). The expected MW for the protein including the His-tag is ~20 kD. B) Digestion of purified Tp2 with Enterokinase using two different incubation times. Samples with Enterokinase (+) are compared to those incubated for the same time but without the enzyme (−). Molecular weights for the protein with and without His-tag are indicated by arrows. C) One dimensional ^1^H NMR spectrum of the monomer fraction of purified Tp2. Note the sharp peaks at either end of the spectrum. D) One dimensional ^1^H NMR spectrum of the dimer fraction. E) NMR spectra from the monomer and dimer fractions overlayed with a focus on the regions with resolved peaks. F) Temperature series of far UV CD spectra of the monomer protein fraction without DTT treatment. Spectra were recorded from 6 to 95°C, indicated by colour gradient from blue to red. The black line indicates the wavelength at which the melting curve was extracted. Minima at 208 and 223 are indicative of a substantial content of secondary structure (α-helix: 14.4%, b-strand: 28.2%, turn: 14.6%, random coil: 42.8%). G) Series of CD spectra recorded for the monomer fraction after addition of DTT. The spectrum is much weaker, and the minima have slightly shifted (205 and 220nm) indicative of a loss of structure. In addition, the same temperature curve from 6 to 95°C causes only a very small change in the CD signal. H) Near UV CD spectrum of the Tp2 monomer fraction without (solid lines) and with DTT (dashed lines). The spectrum in the absence of DTT shows clear negative peaks for the aromatic side chains indicative of the existence of a hydrophobic core while the sample with DTT has no signal at all suggesting the absence of a hydrophobic core.

Monomer and dimer fractions purified using preparative SEC were further analysed by 1D NMR spectroscopy. The spectrum of the monomer fraction contains numerous sharp and well resolved resonance lines in both the high- and low-field regions of the spectrum indicating a folded protein (Fig. 5C). In contrast, the dimer fraction barely contains resolved, visible peaks in the high- and low-field regions of the spectrum, indicative of an unfolded protein (Fig. 5D and 5E). The role of disulphide bonds in the stability of the monomer structure was explored by adding 10 mM of the reducing agent DTT. Significant changes became apparent only after 2h in both high- and low-field ends of the spectrum while full unfolding of the protein was observed after four days (Suppl. Fig. 3E). The NMR properties of the monomer fraction were further explored using a ^15^N labelled sample. The ^1^H-^15^N Heteronuclear Single Quantum Correlation (HSQC) experiment gave an excellent spectrum with about 190 peaks, very close to the number expected for a protein of about 180 amino acids (the protein still had the his-tag attached) (Suppl. Fig. 3C). The peak dispersion is excellent with only a small number of more intense peaks overlapping in the centre.

Circular dichroism (CD) spectra were recorded to determine the overall content of secondary structure and the thermal stability of the monomer fraction. DTT treatment was applied to explore the effect of disulphide bonds on protein structure and stability. The CD spectrum at RT in the absence of DTT clearly belongs to a well folded protein with minima (208 and 223 nm) close to those of α-helices and ß-sheets (Fig. 5F). Best Secondary Structure (BeStSel) ^49^ analysis of the spectrum predicted more than 50% of defined regular secondary structure distributed almost equally between a-helix, ß-sheet and turns. Addition of DTT reduced the intensity of the CD signal to about 20% and caused small but significant shifts of the minima to 205 and 220 nm indicative of a less folded protein (Fig 5G). The near UV CD spectrum of the protein without DTT showed significant peaks which disappeared completely upon addition of DTT (Fig 5H). Only the melting curve of Tp2 without DTT could be analysed giving a melting temperature of 77°C. However, the melting temperature is only a rough estimate because the unfolding does not appear to be completed at approx. 95°C (Suppl. Fig. 3D). It was concluded that the correct protein was synthesised and that at least part of it exhibited a stable and well-defined structure.

### Oral administration of yeast expressing Tp2 induces a antigen specific antibody response in mice

In advance of an immunisation and challenge experiment in cattle, we sought to determine whether oral administration of yeast expressing Tp2 on its surface would induce an immune response in mice. Oral administration of yeast expressing Tp2 resulted in an increase in total IgG serum levels over the study period higher than that observed in the control group that was given yeast expressing the empty plasmid (Fig. 6A). Mice that received yeast expressing native Tp2 had an increased concentration of Tp2 specific IgG antibodies compared to the control group (Fig. 6B). Although these differences weren’t statistically significant, both results indicate that oral administration of yeast expressing specific Theileria antigens increased total as well as induced an antigen specific IgG response.

**Figure 6:**
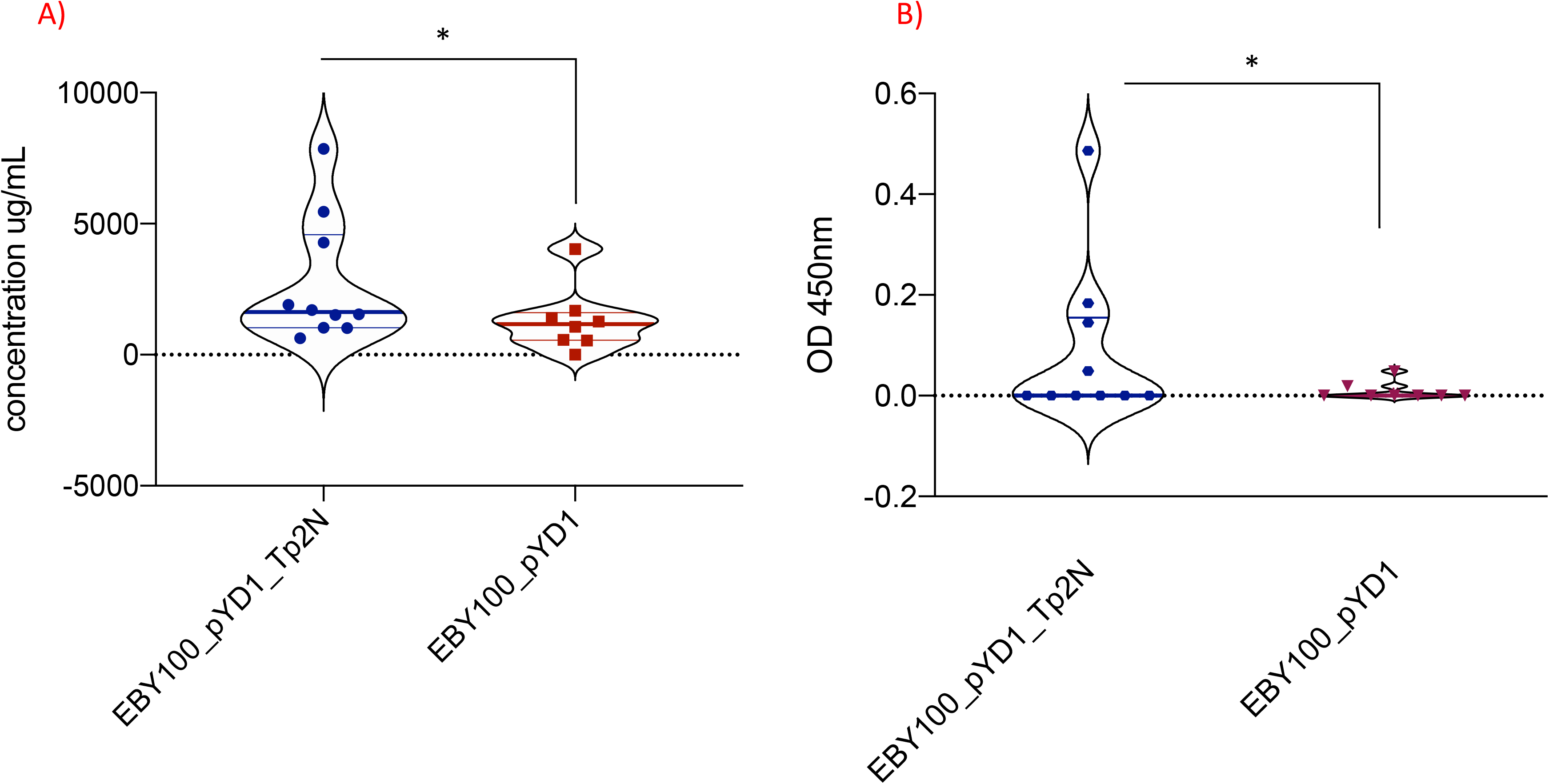
A) Increase in total IgG serum levels over the study period of 90 days. Mice where treated with yeast either expressing native Tp2 (EBY100_pYD1_Tp2N) or carrying an empty plasmid (EBY100_pYD1). Oral administration of yeast expressing Tp2 resulted in a significantly higher increase of total IgG serum levels (p < 0.05) compared to the control group. Where the increase in IgG serum levels produced in response to recombinant yeast was less than background, the concentration is reported as zero. B) Increase in Tp2 specific IgG serum levels over the study period of 90 days. Mice were treated with yeast either expressing native Tp2 (EBY100_pYD1_Tp2N) or carrying an empty plasmid (EBY100_pYD1). Treatment with yeast expressing Tp2 resulted in a significant increase of absorbance (p < 0.05) while no equivalent change could be observed for the control group. Where the increase in absorbance in response to recombinant yeast was less than background, the concentration is reported as zero.

## Discussion

The development of a protective vaccine for ECF is a challenging task. Animals treated with the ITM protocol can remain life-long carriers and can potentially spread vaccine strains into areas previously free from them. It requires a continuous cold chain in liquid nitrogen, co-treatment with oxytetracycline, skilled personnel for delivery and is laborious to produce (reviewed in^50^). Consequently, the development of a subunit vaccine with minimal risks and easier production processes is necessary. Multiple avenues are currently being pursued to achieve this^2,46,51–54^; however, the majority of these approaches appear limited to specific MHC class I haplotypes, or the induction of an antibody response.

We believe that an ideal vaccine platform to protect against *T. parva* infection should induce an early, cross-protective, MHC class I dependent, cytotoxic CD8^+^ T cell response as well as the production of (neutralising) antibodies induced by cross-presentation. Furthermore, it should allow for an affordable large-scale production that is easy to standardise, be transported and administered independently of the farming system from day 1 on (sucklicg calf) and should remain stable at room temperature. These criteria highlighted yeast as an ideal candidate for the delivery of *T. parva* antigens.

Ten proteins expressed by *T. parva* schizonts (Tp1-Tp10) have previously been identified as targets for cytotoxic CD8^+^ T cells ^20,47^, and distinct CTL epitopes were identified in eight of them ^47,55^. In the present study, five antigens (Tp1, Tp2, Tp9, Tp10 and N36) were selected based on the number of CTL epitopes, sequence diversity and presentation by MHC class I molecules. Of these, Tp1, Tp2 and Tp9 were previously identified as antigens with the highest nucleotide and amino acid diversity ^56^. While both Tp1 and Tp9 possess only one known CTL epitope, Tp2 consists of seven potential epitope regions ^43,55^ at least two of which are presented differently by different MHC class I molecules and elicit separate CD8^+^ T cell responses ^43^. This makes Tp2 particularly attractive as a potential vaccine candidate. In contrast, Tp10 could offer broader cross-protection in cattle due to a more conserved sequence with only moderate nucleotide diversity ^56^.

Of the five antigens chosen for expression, we were only able to express the full-length native Tp2 and Tp9 genes in EBY100 yeast, resulting in surface expression of the corresponding proteins. In contrast, we were only able to express fragmented forms of Tp1 within yeast and were unable to transform yeast successfully with Tp10 and N36. Compared to Tp2 and Tp9, genes coding for Tp1, Tp10 and N36 have relatively long reading frames (1,629bp, 1,329bp and 1,662bp, respectively), and these may be too large to be successfully expressed using this yeast-expression plasmid. Furthermore, at least in case of Tp1, glycosylation may have contributed to unsuccessful protein expression. Indeed, Western Blot analysis of recombinant full-length Tp1 and its two fragments, Tp1Na and Tp1Nc reinforced potential glycosylation as all three products did run at a molecular weight substantially higher than predicted.

Recombinant yeast expressing heterologous proteins is considered to be a genetically modified organism (GMO) and therefore strictly regulated. However, inactivated GMO’s are no longer considered to be GMO’s. Therefore, prior to immunisation we performed a heat-inactivation step followed by freeze-drying to achieve thermostability. Heat-inactivation also exposes β-glucans on the yeast cell surface and increases immune recognition of *S. cerevisiae* ^57^. This has proven beneficial as orally administered, heat-inactivated yeast showed increased binding to dectin-1 on M-cells and APCs ^58–60^. The result was induction of mucosal sIgA, systemic IgG as well as cell-mediated immune responses ^24,25,29,30,61,62^ with efficient cross-priming ^32^.

Enhanced uptake of recombinant yeast by APCs should accelerate processing and presentation of Tp antigens to cytotoxic CD8^+^ T cells by MHC class I molecules and to CD4^+^ T helper cells by MHC class II proteins. However, downstream inactivation and freeze-drying of yeast also reduce recombinant protein expression (Dowling and Werling, unpublished data) leading to a potential loss of immunity towards the antigens.

Nevertheless, *ex vivo* exposure of bovine PBMCs to yeast expressing Tp1Nc, native Tp2 and Tp9 successfully elicited IFNγ responses. The absence of any IL-2 production might suggest that out of the T cell population mainly CD8^+^ T cells contributed to the IFNγ response, as IL-2 has been shown to not be necessary for the early induction of a CTL response ^63^. As Theileria specific CD8^+^ T cell mediated immunity is restricted by MHC class I proteins each determined by individual class I genes within the MHC haplotypes present in each animal, MHC class I incompatibility is likely to cause differential antigen recognition by the three animals analysed^47^. Interestingly, IFNγ reactions of enriched cytotoxic CD8^+^ T cells to recombinant yeast were different compared to PBMCs from the same animals. Previous work showed increased IFNγ production from bovine PBMCs towards recombinant full-length Tp9 expressed in HEK293T cells compared to the control ^46^. Depletion of specific T lymphocyte subsets from PBMCs highlighted CD4^+^ rather than CD8^+^ T cells as major source of IFNγ. Thus, it is likely that the change in cell composition from FluoroSpot assay to ELISA analysis could have caused the differential IFNγ responses due to a reduction of CD4^+^ T cells. Additional stimulation of antigen specific CD4^+^ T cells was previously shown to be essential to induce lysis of Tp-infected cells *in vitro* ^16^. In this study, the authors demonstrated that target cell lysis by CTLs required CD4^+^ T cell derived cytokines, while activation of naïve CTLs only occurred after close contact to immune CD4^+^ T cells. Thus, it can be argued that a successful vaccine against *T. parva* schizonts needs to prime both CD8^+^ and CD4^+^ T cells ^16^.

The MHC class I region in cattle is characterised by haplotypes which vary in the number of classical MHC class I genes as well as high levels of allelic diversity in each {Hammond, 2012 #637}. The range of MHC class I diversity was different in both animals studied here, one with a broad range of diversity at four of the predicted classical class I genes and one with a more restricted range of alleles. Differences between these animals in their cellular response to individual antigens does provide additional support to the hypothesis that different MHC class I haplotypes are associated with variation in immune responses towards certain Tp antigens.

Expression of Tp2 in standard *E. coli* strains produced surprisingly high levels of soluble and apparently folded Tp2 protein without the need for assistance with the correct formation of disulphide bonds. The protein contains twelve cysteine residues predicted to form six disulphide bonds. Suggestions for the precise disulphide pattern vary between different prediction programmes, possibly due to the low level of sequence conservation. Both NMR and CD analysis show some evidence for ß-structure which agrees well with known structures of disulphide rich proteins ^64–67^. Although the formation of disulphide bonds in the reducing *E. coli* cytosol is unexpected, evidence clearly supports disulphide bond formation in a variety of proteins in reducing environments ^68^. The existence of a number of different size oligomers suggests that correct formation of disulphide bonds only occurs in about 25% of total synthesised protein. The remaining 75% form a range of higher order oligomers potentially through a disulphide enhanced domain swapping mechanism ^69^. Both NMR and CD spectra indicate the importance of disulphide bonds in maintaining the protein structure as it unfolded after addition of DTT, albeit with a very slow kinetics at room temperature (Suppl. Fig. 3E). This underlines a high level of stability, in agreement with the CD melting curves. It is at present not sure that the bacterially expressed protein has assumed its native structure. However, the excellent quality of the 1D and 2D NMR spectra, the high content of secondary structure, the resistance to thermal unfolding as well as the contribution of the disulphide bridges to the overall stability provide substantial indirect evidence that the protein is folded correctly. These data show that functional and well folded Tp2 can be expressed recombinantly in bacteria which might open interesting avenues for treatment of ECF. The excellent quality of the spectra would even suggest that protein observed NMR screening could be used in the future for small molecule drug development ^70^.

In the absence of suitable small animal challenge models, investigations of the recombinant yeast *in vitro* were followed by evaluating its immunogenicity after oral administration *in vivo*. Yeast itself already possesses strong adjuvant properties as it is recognized by pathogen recognition receptors such as TLR2, TLR4, Dectin-1 as well as DC-SIGN and results in activation of the NF-kB pathway ^59^. Therefore, it was not surprising that we identified a rise in total IgG serum levels in both treatment and control group over the study period of 90 days (Fig. 6A). However, individual mice treated with yeast extracellularly expressing native Tp2 showed an increase in total IgG serum levels higher than that of the control group. Bacterially expressed Tp2 protein was subsequently used in the same setup to further confirm the specificity of the reaction. As expected, oral administration of yeast transfected with the empty plasmid did not increase Tp2-specific IgG serum levels over the study period of 90 days (Fig. 6B). In contrast, mice treated with yeast extracellularly expressing native Tp2 did develop IgG antibodies detectable with purified Tp2.

While our results will require subsequent challenge experiments to verify induction of protection in cattle, these proof of concept data indicate that oral application of yeast expressing Theileria antigens could provide a cheap and easy vaccination platform for sub-Saharan Africa. The freeze-dried yeast, which is stable for many months (unpublished data) would not rely on the presence of uninterrupted cooling chains, could provide a platform to deliver multiple antigen combinations, which could be provided from birth as part of normal milk-replacement supplements. Evaluation of antigen specific cellular immune responses, especially cytotoxic CD8^+^ T cell immunity, in cows will further contribute to the development of a yeast-based vaccine for ECF.

## Supporting information

Genetic subunits of native Tp1 (Tp1N) expressed in pYD1

Cytotoxic activity of CD4+ and CD8+ T lymphocytes

Additional data supporting the synthesis and characterisation of Tp2

Detection of Tp2 expressed by E. coli BL21 Star

## Acknowledgment

This work was funded by the Bill and Melinda Gates Foundation (BMGF) and the Department for International Development (DFID) of the United Kingdom [OPP1078791]. We would like to give our special thanks to Prof. Simon Draper [University of Oxford, Jenner Institute] for his valued comments and suggestions during the planning of this study. KB receives funding from the European Union’s Horizon 2020 research and innovation programme under grant agreement no. 731014KB (VetBioNet) and acknowledges the support received from the Scottish Government’s strategic research programme. We would also like to express our great appreciation to the RVC farm staff who devoted a lot of effort for the animal care and in making sure that the animal studies involving cattle went as planned.

## Author contributions

S.G., D.N, K.T., A.H. D.W.: designed and supervised animal experiments, supervised the MHC class I typing, tested antigen expression, conducted ELISA, ELISpot, flow cytometry assays, performed CTL assays, analysed all the data, generated the figures, and wrote the manuscript; K.B.: performed MHC class I typing and analysed data; J.K.: generated Tp2 vector; M.P. generated purified Tp2 protein and performed biophysical studies; J.K.: analysed murine serum samples and developed Tp2 ELISA. S.D. provided input in the experimental set-up; D.W.: conceptualized and designed experiments, supervised immunology experiments, provided inputs to analyses of data; S.G., A.H., J.K and D.W. wrote manuscript; All authors agreed on manuscript

