## Supplementary material for "Development of a yeast-based vaccine for *Theileria parva* infection in cattle": Genetic subunits of native Tp1 (Tp1N) expressed in pYD1

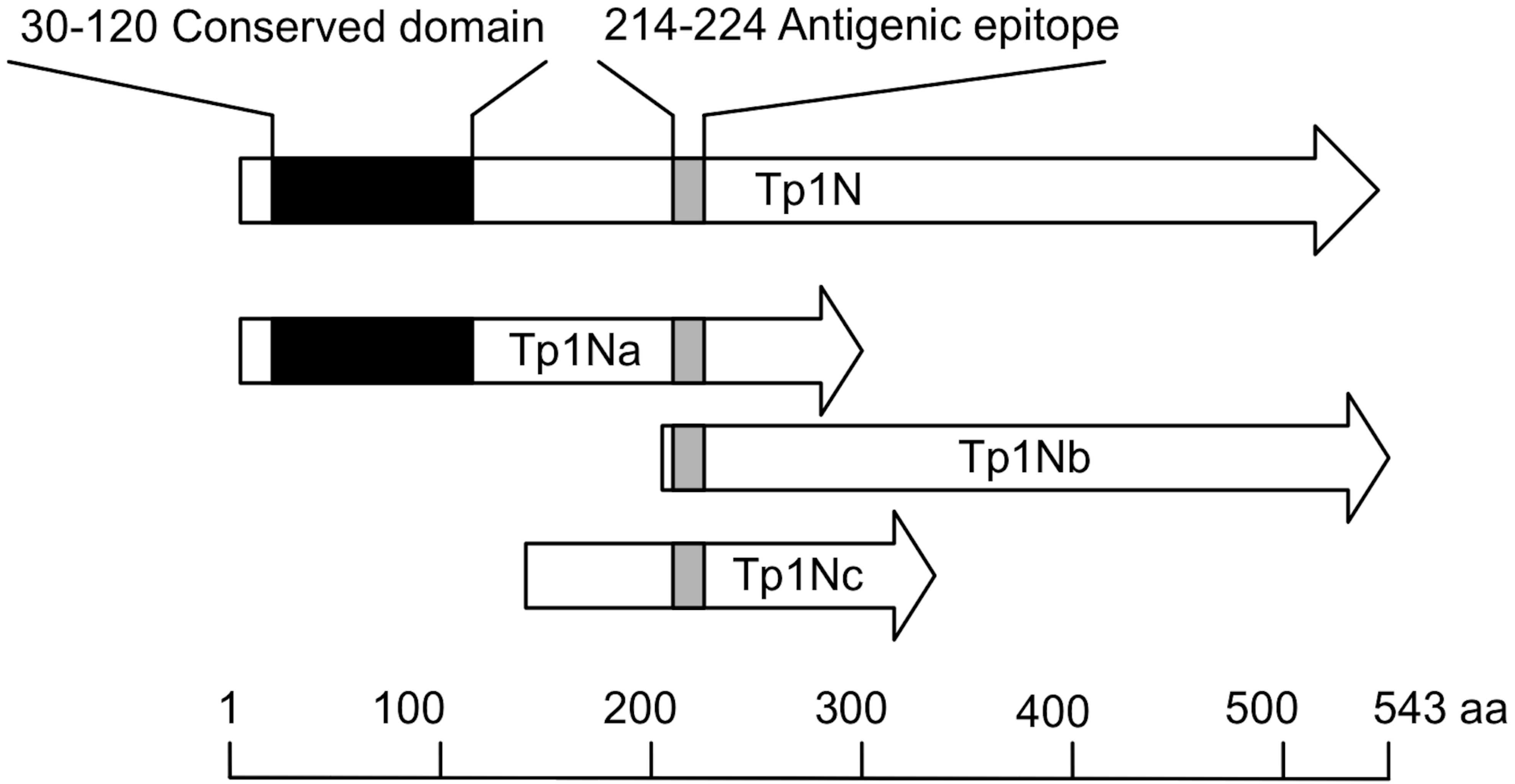


**Supplementary Figure 1 : Genetic subunits of native Tp1 (Tp1N) expressed in pYD1. Known features of native Tp1 are indicated and the antigenic CD8$^{+}$ T cell epitope is present in all subunits.**
