## Supplementary material for "Development of a yeast-based vaccine for *Theileria parva* infection in cattle": Cytotoxic activity of CD4+ and CD8+ T lymphocytes

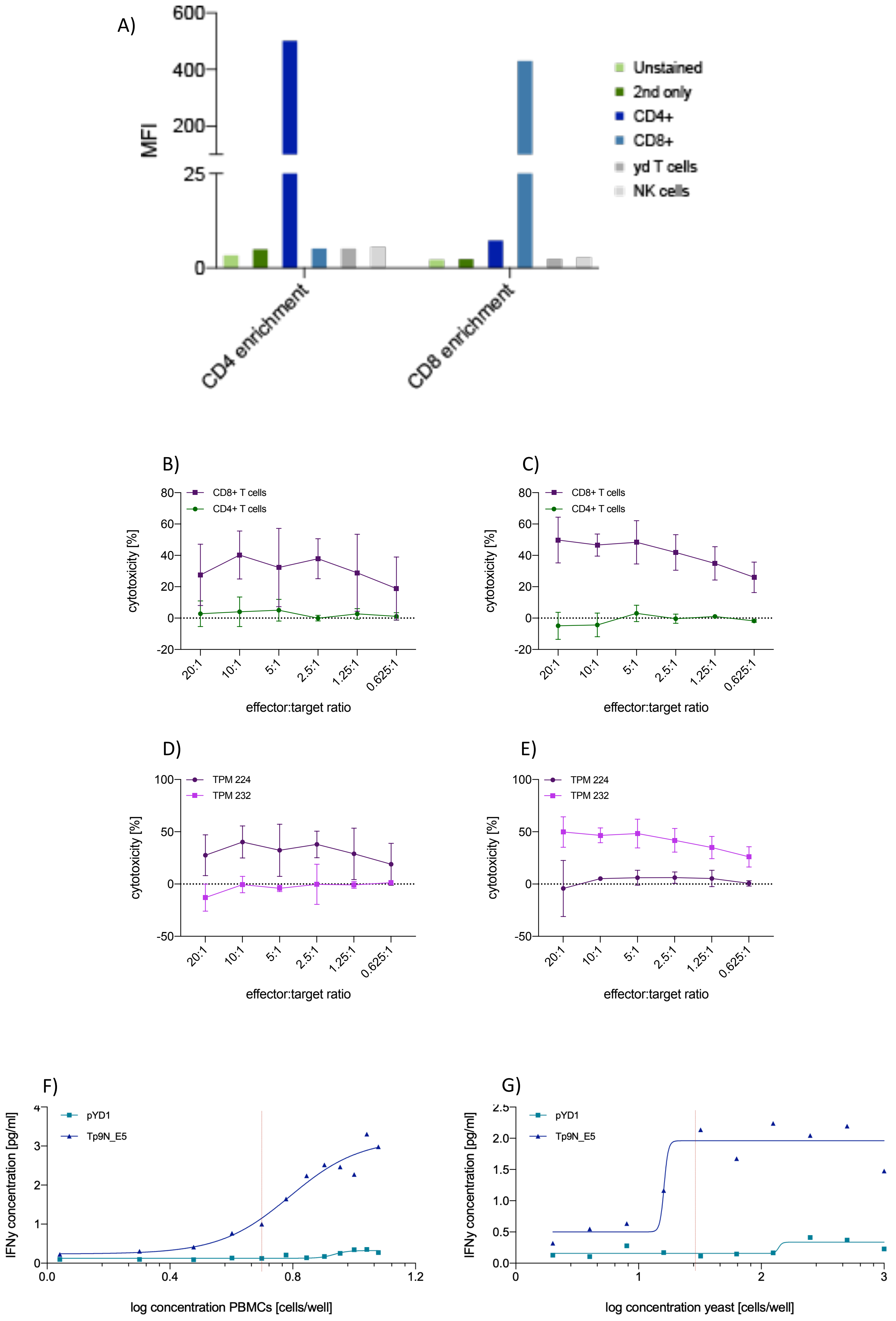


Suppl. Fig. 2: Cytotoxic activity of $CD4^+$ and $CD8^+$ T lymphocytes. A) Median fluorescence intensity of bovine PBMCs enriched for either $CD4^+$ T helper cells or $CD8^+$ T lymphocytes by complement fixation. B-E: Cytotoxic activity of $CD4^+$ T helper cells compared to $CD8^+$ T lymphocytes from animals B) 224 and C) 232 towards TpM infected cells at various effector:target ratios. $CD8^+$ T cells lysed TpM infected targets while $CD4^+$ T helper cells did not respond. However, the lytic activity of $CD8^+$ T lymphocytes from cow 224 (D) and 232 (E) was restricted to target cells originating from the same animal. F-G: IFNγ production of effector cells and APCs (cow 224) towards EBY100 yeast with either empty pYD1 plasmid (green) or pYD1_Tp9N (blue). APC and yeast concentrations were transformed to log scale for subsequent non-linear regression analysis. F) $2.5x10^4$ $CD8^+$ T lymphocytes were incubated with $5x10^4$ yeast cells and increasing numbers of APCs. The red line indicates the number of APCs selected for IFNγ ELISA. G) $2.5x10^4$ $CD8^+$ T cells were incubated with $5x10^3$ APCs and increasing numbers of yeast cells. The red line indicates the number of yeast cells selected for the IFNγ ELISA.
