## Additional data supporting the synthesis and characterisation of Tp2 for "Development of a yeast-based vaccine for *Theileria parva* infection in cattle"

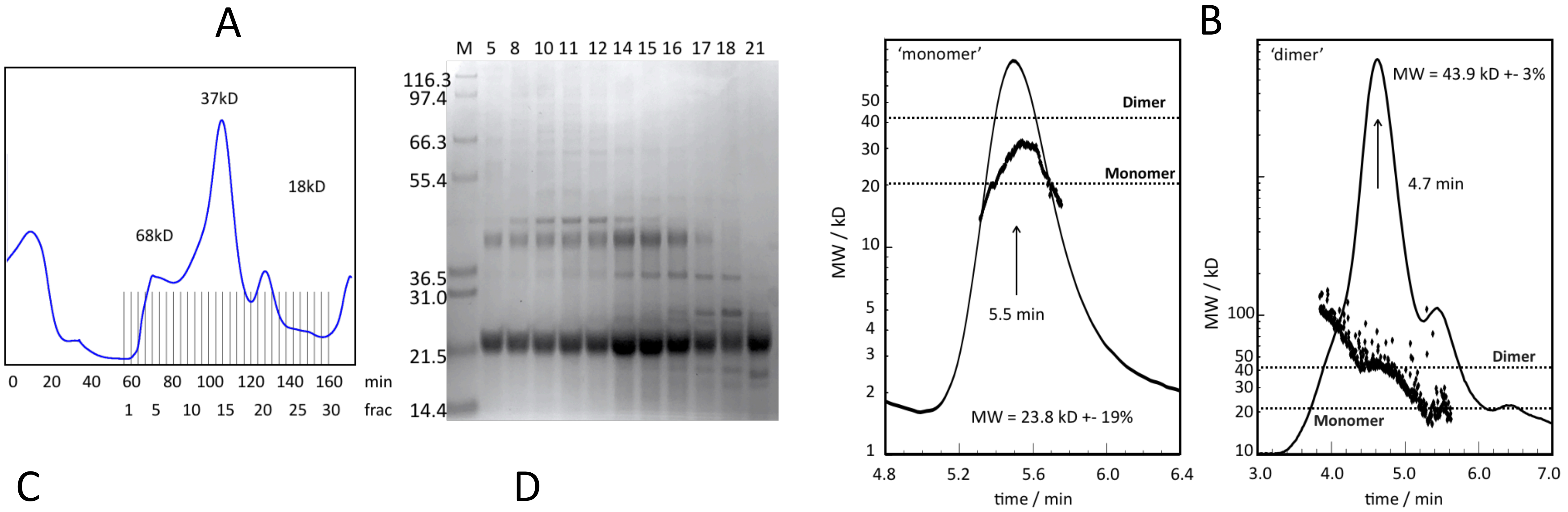


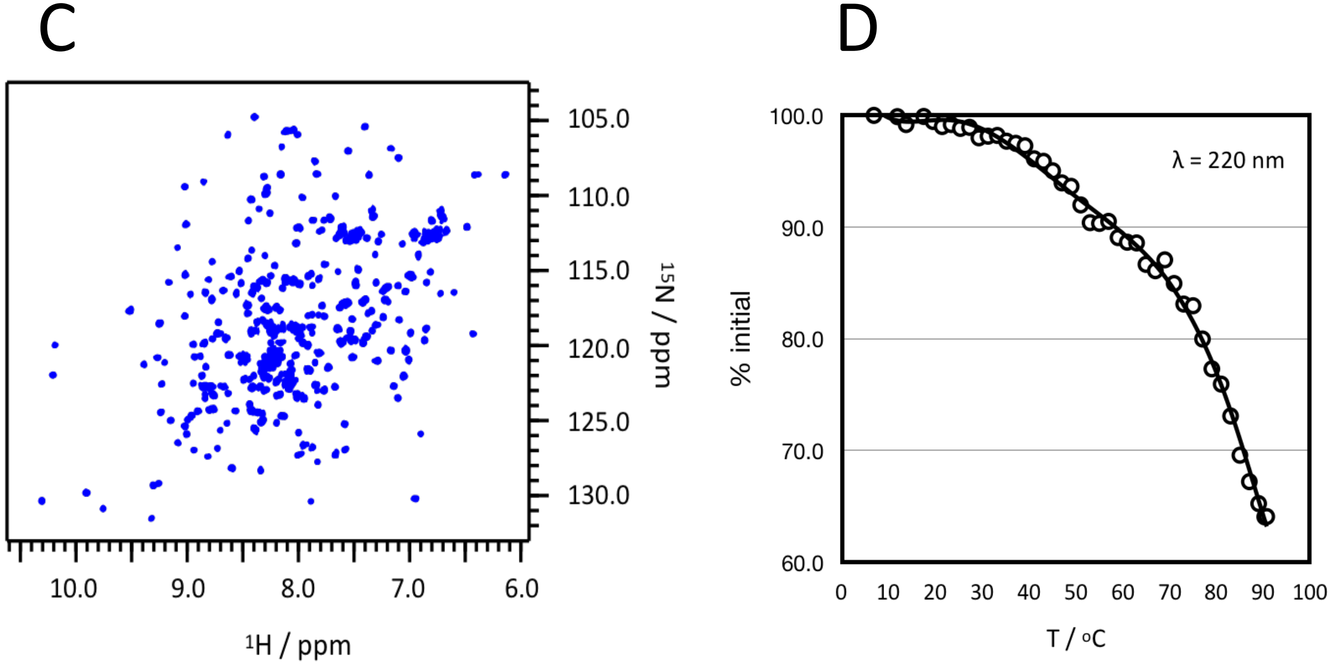


Supplementary Figure 3: Additional data supporting the synthesis and characterisation of Tp2 . A) SEC purification of Tp2. Apparent MW based on column calibration are indicated in the chromatogram. Selected fractions are analysed on SDS-PAGE. B) SEC-MALS of pooled monomer and dimer fractions. Absorption chromatogram is shown as continuous black line, light scattering is shown in black dots. Monomer and dimer molecular weights are indicated for guidance. Peak positions are shown by arrows and fitted molecular weights from light scattering are shown. C) 1H-15N HSQC spectrum of monomer Tp2. The very good peak dispersion and uniform linewidths indicate a well folded protein. D) Thermal unfolding curve of the Tp2 monomer taken from Fig 5F.

E

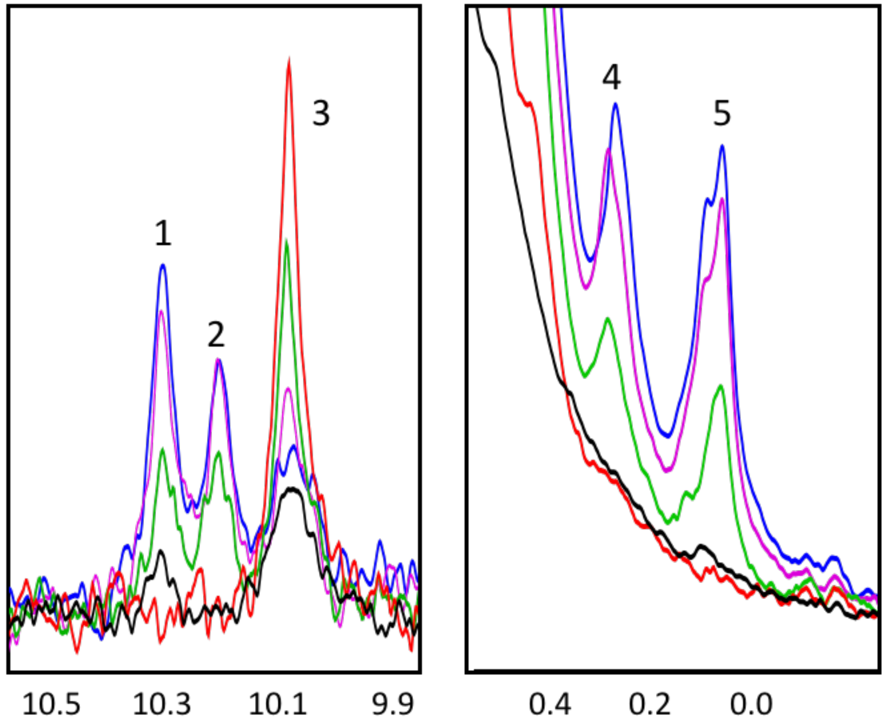


Supplementary Figure 3, continued: E) Effects of DTT on the conformation of the Tp2 monomer. Well resolved peaks from the low field (1-3) and high field (4 & 5) region of the spectrum are indicative of a folded conformation. Spectra are: Blue: Tp2 monomer, no DTT; Magenta: Tp2 monomer + 10 mM DTT, 10 mins; Green: Tp2 monomer + 10 mM DTT, 2h; Red: Tp2 monomer + 10 mM DTT, 4 days; Black: 1D spectrum of Tp2 dimer as reference. Addition of DTT to the Tp2 monomer leads to a disappearance of well resolved peaks indicative of a folded protein. Exception is peak 3 which actually gets stronger and is also seen in the un/misfolded Tp2 dimer. This is very likely the indol resonance of one of the tryptophanes in Tp2 which appears in this area even in unfolded proteins.
