## Supplementary material for "Development of a yeast-based vaccine for *Theileria parva* infection in cattle": Detection of Tp2 expressed by E. coli BL21 Star

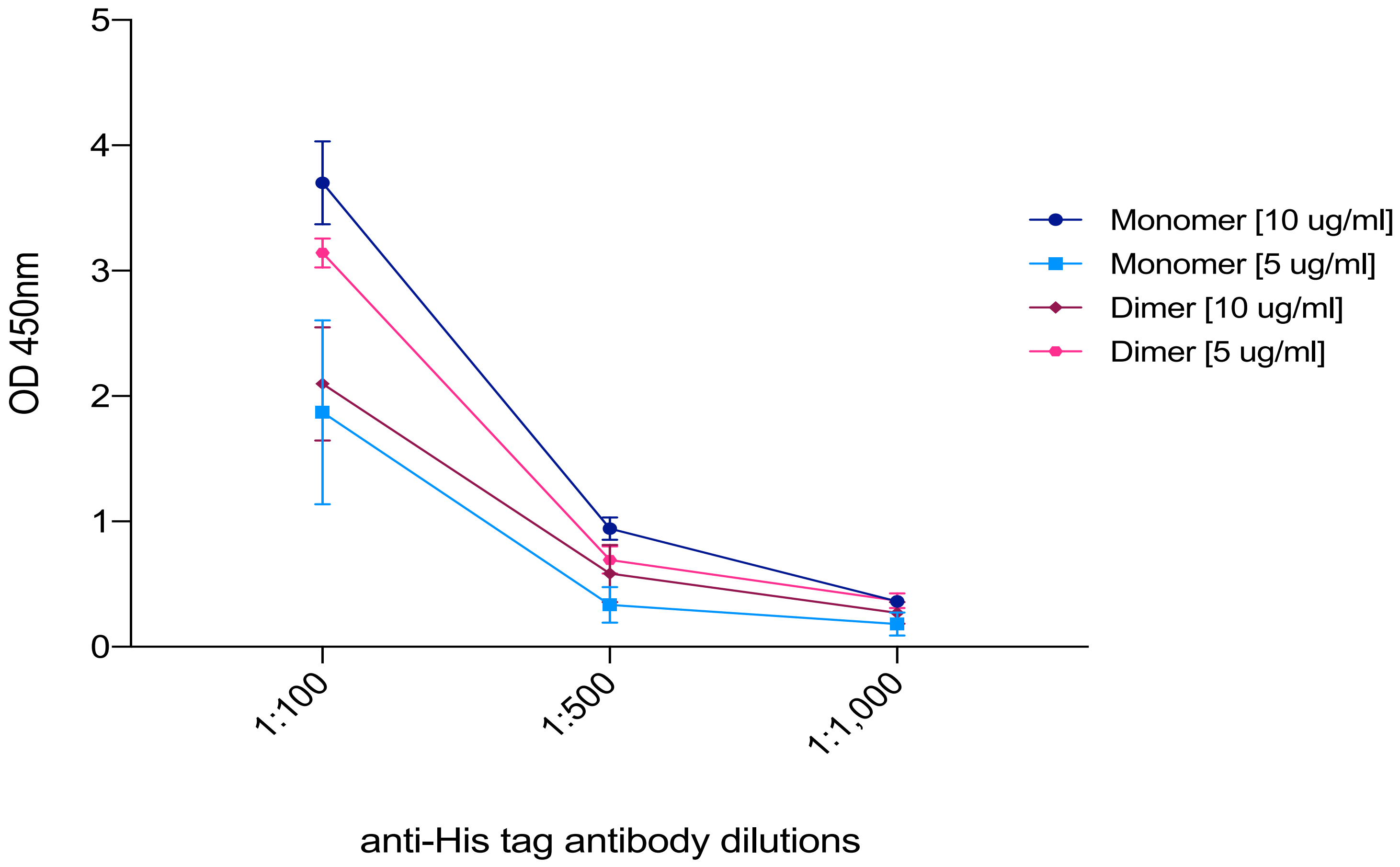


Supplementary Figure 4: Detection of Tp2 expressed by *E. coli* BL21 Star on Nunc MaxiSorp™ Flat bottom 96-well ELISA plates after overnight incubation at 4°C. Binding of both monomer and dimer fractions (10 µg $ml^{-1}$ and 5 µg $ml^{-1}$) was evaluated with three dilutions of an anti-Histidine tag monoclonal antibody. Each fraction was detected by the antibody in a dose-dependent manner confirming proper binding of the protein to the plate. However, lower concentrations of dimer resulted in a stronger signal compared to higher concentrations. Consequently, the monomer was selected for further experiments. The results displayed represent one of two independent experiments.
